# Explainable AI identifies H3K18ac as a new marker of active enhancers

**DOI:** 10.64898/2026.06.09.731088

**Authors:** Khizra Maqsood, Duarte Polvora Brandao, Jareth Wolfe, Bhavana Kayyar, Catalin-George Clinciu, Roxana Andreea Bosnea, Olivia A. Grant, Fanny Boulet, Christopher G. Bell, Gabriella Ficz, Madapura M. Pradeepa, Hani Hagras, Nicolae Radu Zabet

**Author notes:** Co-first authors (Khizra Maqsood, Duarte Polvora Brandao).

## Abstract

Enhancers are non-coding regions of DNA that regulate gene transcription, yet the mechanisms underlying enhancer activity remain incompletely understood. Despite extensive experimental and computational efforts, we still lack accurate enhancer maps in many human cells, tissues and disease contexts. Here, we developed several Artificial Intelligence (AI) models (Convolutional Neural Networks (CNN), XGBoost, Logistic Regression (LR) and an eXplainable Artificial Intelligence type2 Fuzzy Logic based System (type2-FLS)) to predict enhancers across different human and mouse cell lines. While all models display high accuracy in the cell lines they were trained on, our results confirmed that type2-FLS, and, partially, CNN, LR and XGBoost perform consistently well in cell lines unseen during training, supporting the generalisation of the models. Most importantly, type2-FLS identified H3K18ac as an important enhancer mark along with many novel putative enhancers, which display the same epigenetic signatures as experimentally identified ones. We have validated some of these novel enhancers by both global epigenetic perturbations and directed enhancer epigenetic rewriting (CRISPRi). Interestingly, seven epigenetic marks in humans and five in mouse are sufficient to annotate enhancers without losing accuracy. Overall, we have deciphered the epigenetic code of mammalian enhancers and annotated enhancers in multiple human and mouse cell lines.

## Introduction

Enhancers are non-coding regions in the genome that control the levels and timing of gene expression in various cell types and tissues ^1^. Sequences with enhancer activity harbour a high density of DNA motifs recognised by activating transcription factors and architectural proteins ^2^. However, there is no general sequence code for enhancers, and a single enhancer can control multiple genes ^3^. Each gene can also have multiple enhancers that control it in different tissues or cell types, or even in the same tissue or cell type ^2^. Furthermore, unlike promoters, which are located proximally to the transcription start site, the position of an enhancer relative to its target gene is highly variable and can occur upstream, downstream or within introns ^4^. In addition to not having a specific location in the genome and no general sequence code, enhancers can also be active only in certain spatial, temporal, or environmental conditions ^5^. Altogether, this makes enhancer annotation challenging.

Despite the lack of a sequence code, there are specific epigenetic signatures that are associated with enhancers, including post-translational modifications of histones. The two most common histone modifications linked with enhancer identification and activation are the mono-methylation of lysine 4 and the acetylation of lysine 27 of histone H3 (H3K4me1 and H3K27ac, respectively) ^6^. These two marks have been used to identify enhancers and to separate active enhancers from poised ones ^7^. However, recent works showed that they alone do not indicate enhancer activity, and several other marks have been associated with active enhancers, such as H3K9ac, H3K14ac, H3K18ac, H3K23ac, H3K122ac and H4K16ac ^8–14^. This could potentially explain the relatively reduced overlap between enhancers identified by using the regions showing enrichment of both H3K27ac and H3K4me1 and enhancers identified by other methods (including FANTOM enhancers, GRO-CAP, P300 ChIP-seq, etc.)^15^.

Furthermore, studies in Drosophila and mice have shown that H3K27ac may be dispensable for gene activation, despite its strong correlation with active transcription ^16–18^. H3K27ac is deposited by the lysine acetyltransferases CBP and p300, which are also responsible for the acetylation of the neighbouring K18 site (while both K14 and K23 are unaffected, showing its precise targeting) and multiple other protein targets, which may lead to confounding effects ^19,20^. Thus, although H3K27ac is the most common mark used to annotate enhancers (identifying genomic loci with enhancer activity based on genomic data), it represents only a part of the full repertoire of histone modifications in these regulatory regions ^21,22^.

H3K18ac has been previously reported to be enriched at enhancers ^22–24^. Previous work showed that SIRT7 mediates repression of L1 elements and upon depletion, H3K18ac is accumulated at these L1 elements leading to their expression ^25^. Another study has shown that p300 inhibition (which would result in loss of H3K27ac and H3K18ac) was linked with silencing of ERVK elements in 8-cell embryos ^26^. Nevertheless, inhibition of p300 by siRNA was large and lead to downregulation of H3K27ac so it was not possible to disentangle the effects of H3K27ac and H3K18ac. Moreover, recent works have shown that H3K18ac promotes accessibility to H3K4 of methyl writers and readers, indicating a mechanism of crosstalk between H3K18ac and H3K4 methylation in transcriptional regulation ^27,28^. Overall, there is evidence that H3K18ac could be a functional enhancer modification.

Massively Parallel Reporter Assays (MPRAs), including STARR-seq, have been successfully used to annotate experimentally active enhancers ^3,29^. However, a significant challenge remains beca[NO_PRINTED_FORM]use MPRAs are technically complex and resource-intensive for genome-wide applications, which is in contrast with generating genome-wide epigenetic profiles that is relatively scalable. Consequently, developing computational approaches to predict enhancer activity as measured by MPRAs/STARR-seq from epigenetic data is a scalable strategy for improving enhancer annotation across diverse cells, tissues and disease states.

Recently, computation tools, specifically machine and deep learning tools, have been used to annotate enhancers genome-wide ^30^. These tools used histone modification profiles and high-throughput sequencing assay data for training and can predict enhancers genome-wide. Machine learning methods, such as Convolution Neural Networks ^31^, Hidden Markov Models ^32^, Random Forest ^33^, Dynamic Bayesian Network ^34^, and Support Vector Machine^35^, have been developed and can annotate enhancers with different degrees of accuracy, but require very large amounts of data as training and validation sets. Thus, predictions could suffer from overfitting and do not generalise to other cell types, tissues or disease conditions when the training set is not large enough ^36^ and can have difficulties at predicting cell type specific putative regulatory regions ^37^. Most importantly, opaque box models can be interpreted, for example using Shapley Additive exPlanations (SHAP) ^38^, which can rank individual features based on their importances and occasionally reveal some combinatorial interactions ^39^. However, they frequently fail to systematically articulate the conditional, multi-feature logic that governs the enhancers underlying mechanism; for example, where a region is annotated as an enhancer if H3K27ac and H3K4me1 are both enriched and H3K4me3 is depleted. This is where rule-based explainable AI (XAI) models can help by generating natural language IF/THEN rules as a classification algorithm based on type2 Fuzzy Logic Systems (type2-FLS) optimised via multi objective multi constraint genetic algorithm (to maximise explainability via generating the smallest set of short IF-THEN rules that have the possible highest accuracy, thus, allowing to generate fully explainable models that are close in accuracy to black box models) ^40–42^. Type2-FLS helps us model how features interact in high-dimensional biological data, handling some major drawbacks of post-hoc explainability methods like SHAP ^38^ and LIME ^43^. Previously, we showed that type2-FLS rule-based explainable artificial intelligence models can predict the location of known enhancers in *Drosophila* and provide insight into the combinatorial histone modification code underpinning enhancer activity ^22^. ^44,45,46^ Based on that, in this manuscript, we apply an explainable AI method (type2-FLS) and three other ML methods (Convolution Neural Networks, XGBoost and Logistic Regression) to human and mouse genomic data to annotate enhancer regions using only epigenetics data. By doing so, we demonstrate that our models not only predict enhancer activity with high accuracy but also provide clear, biologically interpretable rules. Our results identifies H3K18ac as a critical, under-appreciated marker for enhancer annotation, capable of defining a set of putative novel enhancers that lack the classical H3K27ac signature.

## Results

### ML and eXplainable AI predict enhancers from epigenetic data and identify a set of novel enhancers

We employed an explainable AI method (type2-FLS) and three other machine learning methods to predict enhancers (as predicted by STARR-seq) using histone modification genome-wide profiles (e.g., ChIP-seq), open chromatin data (e.g., ATAC-seq or DNaseI-seq), and methylation data (e.g., WGBS). In particular, we trained four Machine Learning and AI models for predicting the locations of enhancers, namely: type2 Fuzzy Logic Systems based XAI (type2-FLS), Logistic Regression (LR), XGBoost and Convolutional Neural Network (CNN). The entire genome was split into 100 bp tiled bins, and the values of each epigenetic dataset within each bin were stored in a matrix (see Figure 1 and *Materials and Methods*). First, we trained and validated the models in H9 human Embryonic Stem cells (hESCs) and all the ML/AI models exhibited robust predictive power (AUC between 0.79 and 0.87; AUCPR between 0.05 and 0.09); see Figure 2A, Figure S1A and Supplementary Table S1.

**Figure 1:**
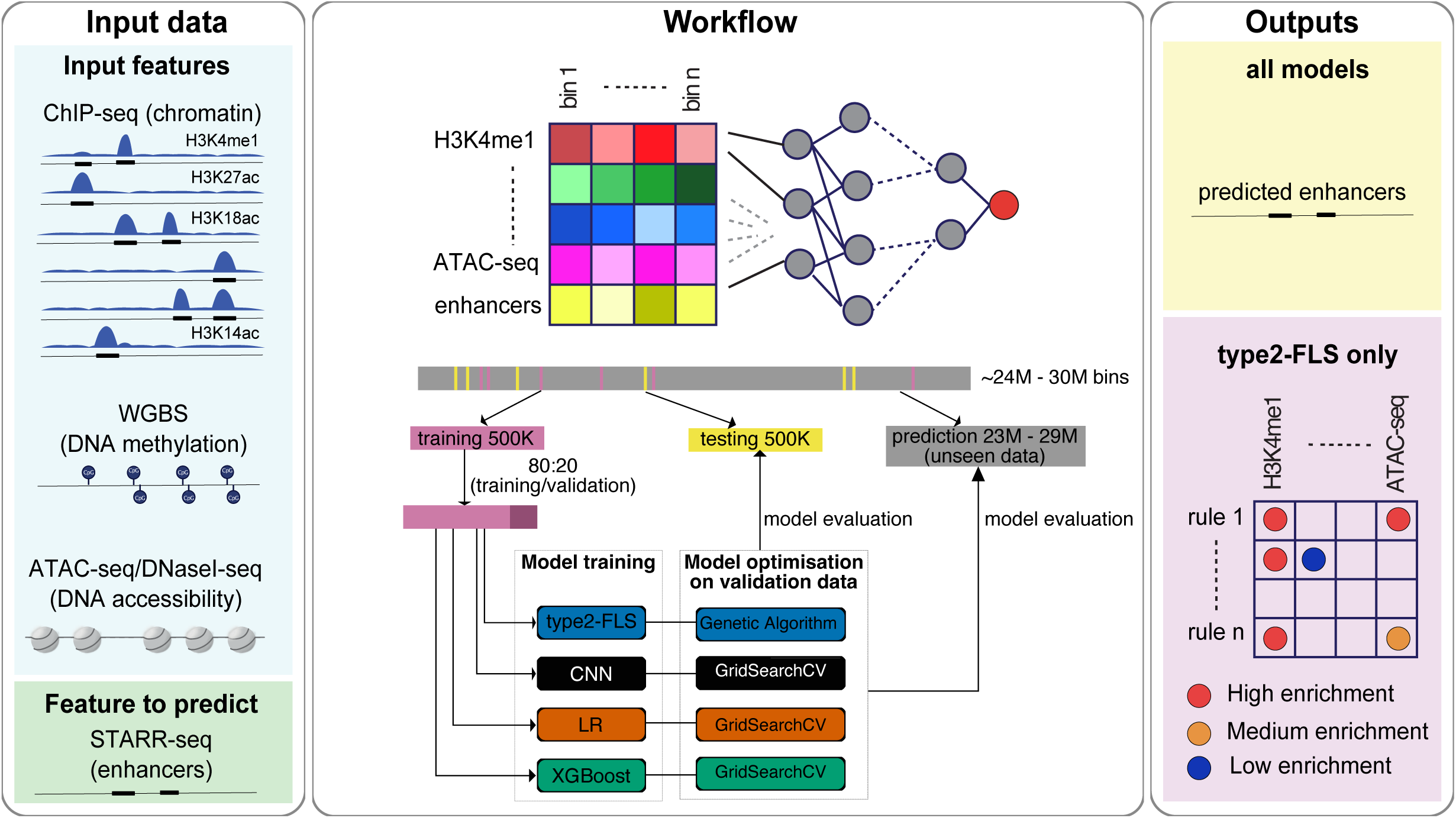
Workflow to annotate enhancers. We start with several epigenetic datasets, including ChIP-seq of histone modifications, ATAC-seq and WGBS and use enhancer annotation from multi-parallel reporter assays, such as STARR-seq. The genome is split into 100bp tilling bins, and the normalised signal in each bin for each epigenetic feature is computed. Then we train several ML/AI models (CNN, LR, XGBoost and type2-FLS) and evaluate the performance of these models on unseen data, which includes other cell types or other organisms. The opaque box models provide only the predictions of enhancers, while type2-FLS also includes the rules used to make the annotation (in the form of IF/THEN rules).

**Figure 2:**
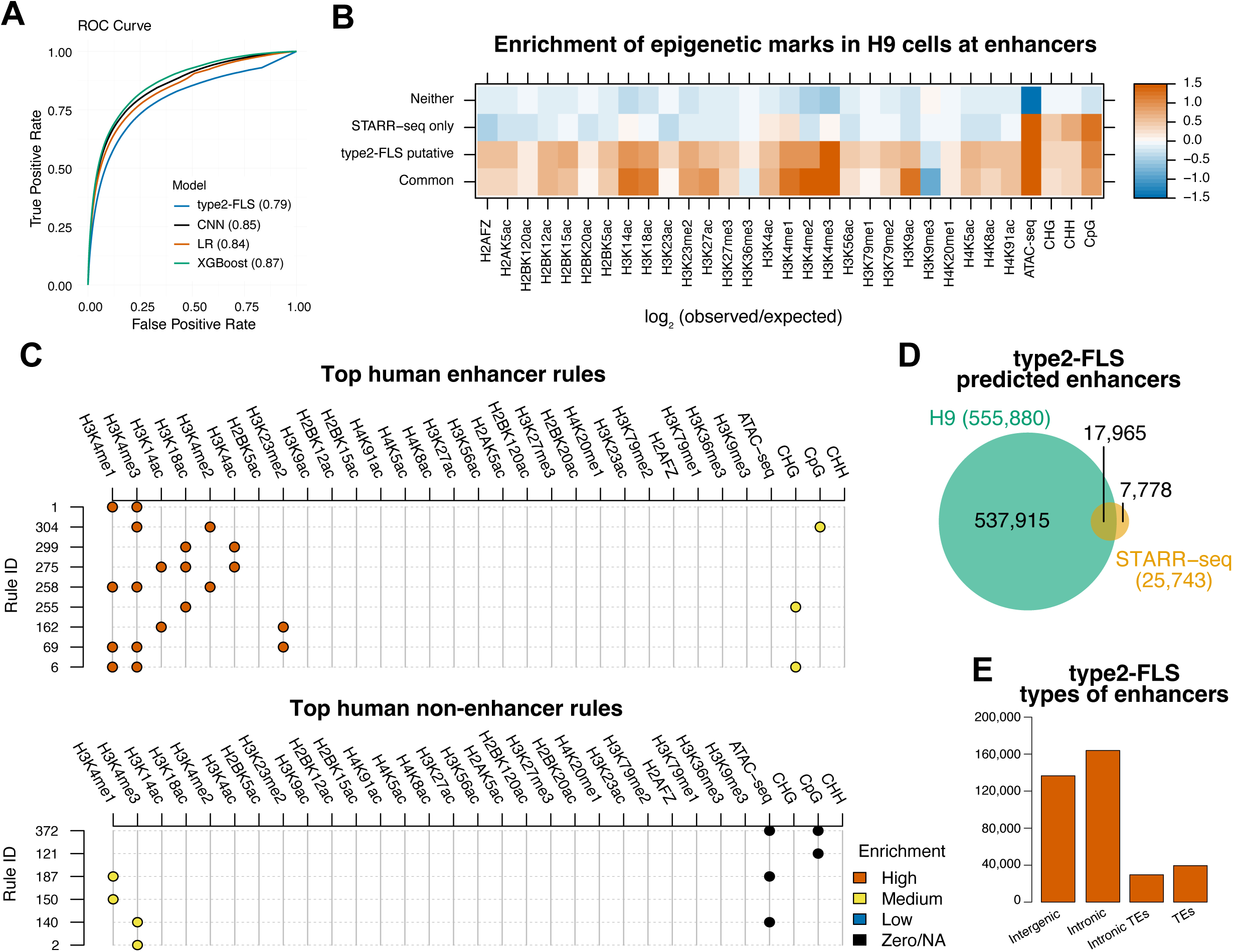
Annotation of enhancers in human ES cells. *(A)* ROC curve and AUC values for four ML/AI models (CNN, LR, XGBoost and type2-FLS) trained on 31 epigenetic features in H9 human ES cells. Training was performed on 500K bins, and the evaluation was performed on all unseen data. *(B)* Enrichment of all 31 epigenetic signals (log_2_ observed/expected) at common enhancers (shared between type2-FLS and STARR-seq), putative enhancers (type2-FLS only), STARR-seq only enhancers and regions that are not classified as enhancers by either of the methods. *(C)* The top rules to classify enhancers (top) and non-enhancers (bottom). Each row represents a rule, and the colour represents the enrichment of the signal (red for high, yellow for medium, blue for low and black for zero/NA). *(D)* Venn diagram showing the overlap between annotated enhancers in H9 cells and STARR-seq enhancers. Annotated enhancers are computed by merging individual bins (see *Materials and Methods*) and selecting non-promoter enhancers of at least 200 bp in length. *(E)* Barplot with the number of annotated enhancers by type2-FLS that are located in intergenic regions, introns, intronic TEs and TEs.

The lower precision and AUCPR for all ML/AI models (Figure S1A) are due to a higher number of regions predicted to be enhancers not identified experimentally by ChIP-STARR-seq. It is also worthwhile noting that only approximately 0.8% of the bins were identified as enhancers in H9 cells by ChIP-STARR-seq. To investigate this further, bins with a probability of at least 0.8 of being an enhancer and within 500 bp of each other were merged into regions that we labelled as predicted enhancers (see *Materials and Methods*). Most of the enhancers fit the expected size of 100bp to 500bp regions, but we also found some larger enhancers (greater than 1 Kb) that could be potential super-enhancers (Figure S1B). For the downstream analysis, we selected predicted enhancers not located within the promoter region (up to 1Kb upstream of TSS) that are at least 200 bp (at least two neighbouring bins). Note that enhancers smaller than 200 bp do exist, but here we wanted to select enhancers where at least two neighbouring bins have a high probability of being enhancers, which ensures that the selected regions have a high overall probability of being enhancers and are not affected by spurious local signals.

Given that we merged enhancers only if the entire region displayed a probability of at least 0.8, there is still the possibility that we are detecting clusters of enhancers. This is indeed the case, and we found that many of the enhancers are within 1-10 *Kb* of each other (Figure S1C). The total number of enhancers (555.9 K) drops to 333K when merging enhancers within 1 *Kb* and to 69K when merging enhancers within 10 *Kb,* independent of the probability of the entire region being an enhancer (Figure S1D). This indicates that we identified small DNA regions (200-400 bp) with enhancer activity, which can drive expression and these small enhancer regions cluster within ∼70K larger genomic regions.

### type2-FLS based XAI model provides explainable rules for the annotation of enhancers

To investigate the signatures of the identified enhancers in human H9 cells (using the H9 31 features trained models), we compared the enrichment of different epigenetic modifications at putative enhancers (identified by type2-FLS only), common enhancers (identified both by type2-FLS and STARR-seq) and enhancers identified by STARR-seq only. Our results showed that putative enhancers and enhancers common between type2-FLS and STARR-seq display similar epigenetic signatures, but the putative enhancers displayed slightly less enrichment for H3K4me1, H3K4me2, H3K14ac, H3K18ac and H3K9ac (Figure 2B). This indicates that the novel putative enhancers might have lower activity and could be missed by STARR-seq. Interestingly, enhancers that are not captured by type2-FLS (STARR-seq only enhancers) have enrichment only for H3K4me1 and ATAC-seq but lack all the other histone modifications, which may suggest that they could function as enhancers in the plasmid-based in vitro reporter assays, but not in the endogenous chromatin context ^47,48^.

Most importantly, our type2-FLS-based XAI model allows us to extract the top rules for predicting regions as enhancers or non-enhancers (Figure 2C and S2). We found H3K4me1, H3K4me2 and H3K4me3 to be the top epigenetic modifications associated with enhancers. ATAC-seq signal has been previously used as a proxy to identify enhancers ^49,50^, but our results show that the signal from ATAC-seq is not sufficiently specific to classify regions as enhancers. Nevertheless, lack of accessibility is a top rule to predict a region as not being an enhancer, i.e., enhancers need to be accessible, but not all accessible regions are enhancers. Although H3K27ac is a canonical enhancer-associated histone modification^51^, we find it enriched only in lower-dominance regulatory patterns of human enhancer (Figure S2); where the dominance is a metric that combines the rule support and its confidence (see *Materials and Methods*) ^52^. Our results are supported by recent work that showed that depletion of H3K27ac in mouse ESCs results in only a few small changes in gene expression ^53,54^. Finally, DNA methylation is usually associated with repression of transposons and genes ^55^. However, we found that a complete lack of DNA methylation is associated with regions that are not enhancers, while intermediary levels of DNA methylation can be predictive of enhancers (see Figure 2C bottom panel), a result that has been reported recently ^56^. Interestingly, we found that H3K18ac and H3K14ac, two epigenetic modifications whose association with enhancers was previously reported ^57,58^ but often neglected, are epigenetic modifications strongly predictive of human enhancers. H3K18ac was also one of the histone modifications most associated with enhancers in *Drosophila*, by our type2-FLS models^22^.

While Opaque Machine Learning models are not explainable, they can be interpreted ^59^, for example, using the SHapley Additive exPlanations (SHAP) method ^38^. We have computed SHAP coefficients for the CNN and XGBoost models and also reported the coefficients from the Logistic Regression. Unsurprisingly, despite the three models being trained on the same dataset, each displays different features as being the most important ones (Figure S3). In particular, CNN identifies H3K79me1, H4K5ac and H2FAZ, while XGBoost identifies H3K4me1, H3K4me3 and CpG methylation as the most important features. In contrast, Logistic Regression identifies H3K4me1, H3K4me3 and ATAC-seq signal as the most important features. Despite all three models performing well (AUC ≥ 0.84) and being trained on the same data, the fact that each of the models has a different set of important features according to the SHAP coefficients indicates that interpretation of these opaque box models, at least with SHAP values, needs to be taken with some caution when using those explanations to assign biological function or mechanism, especially for Deep Learning models. Furthermore, these interpretations cannot account for complex logical scenarios (e.g., combinations of different features) as opposed to type2-FLS.

### Characterisation of the novel putative enhancers

For H9 cells (using the H9 31 features trained models), our type2-FLS-based XAI model recovered most of the STARR-seq annotated enhancers (17.9K out of 25.7K) and predicted around 538K novel enhancers, which we labelled as putative enhancers (Figure 2D). To investigate whether the type2-FLS putative enhancers were previously annotated by different methods, we compared them with Enhancer Atlas 2.0 enhancers and found that only approximately 22.5K of the 538K putative enhancers were previously reported ^60^ (Figure S4A). These putative enhancers are similarly distributed in the genome to the ones identified by both STARR-seq and type2-FLS, with a majority being intronic and intergenic (85%), but also approximately 15-20% being derived from Transposable Elements (TEs), which indicates that TEs-derived enhancers establish the gene regulatory networks and significantly contribute to gene regulation (Figure 2E and Figure S4B-C) as recently reported ^13,61,62^.

Of all intronic and intergenic enhancers (300,551), approximately one-third (97,760) are proximal to a promoter (less than 25 Kb), with another third (103,676) located at large distances from the nearest promoter (greater than 100Kb); Figure S5A. There are negligible differences between the distribution of proximal or distal putative enhancers and enhancers identified by type2-FLS and STARR-seq (Figure S5B). Interestingly, both proximal and distal enhancers can be linked to an expressed gene (Figure S5C), but only distal enhancers are linked to higher-expressed genes (Figure S5D). However, this is the case for STARR-seq enhancers, putative enhancers and even the background, suggesting that 3D contacts will link regulatory regions with highly transcribed genes. This supports the idea that these regions may be transcription factories ^63^.

Interestingly, most type2-FLS proximal intronic and intergenic enhancers (97,760) are novel (96,392), and of these, approximately two-thirds have been labelled by the type2-FLS model as having medium or high signal for H3K18ac. Most importantly, approximately half of those (32,552) do not have enrichment of H3K27ac, but they are enriched for H3K18ac, indicating that H3K18ac has an important role for human ES enhancers.

### Putative enhancers display high levels of nascent transcription

We compared how well each of the four ML/AI models recovered enhancers in H9 cells identified by the other three ML/AI methods (using the H9 31 features trained models) and STARR-seq; see Figure S6A. Type2-FLS identifies most STARR-seq enhancers but also exhibits the highest number of predicted putative enhancers. We selected as common, enhancers detected by both type2-FLS and STARR-seq and split our putative predicted enhancer sets into two groups: *(i)* putative type2-FLS only (enhancers only predicted by type2-FLS that do not overlap with the STARR-seq annotated enhancers) and *(ii)* putative multiple (enhancers predicted by type2-FLS and other ML methods but not by STARR-seq). Furthermore, both groups of putative enhancers displayed genomic distribution similar to those identified by STARR-seq, with a majority being intronic and intergenic (71%), but also approximately 29% being derived from TEs (Figure S6B).

Active enhancers produce eRNAs ^64^ and, thus, display higher level of nascent transcription than genomic background. To account for differences in expected nascent transcription depending on the genomic context, the enhancers were further divided into intergenic, intronic, intronic TEs and TEs. For intronic and intronic TEs enhancers, background levels were computed using randomly selected bins in introns of genes harbouring STARR-seq enhancers to ensure direct comparison within transcriptionally active loci (see *Materials and Methods*). Across all genomic categories, common enhancers and putative multiple enhancers exhibited significantly elevated nascent transcription compared with the background (Figure 3A). Importantly, putative type2-FLS only enhancers located in introns or intronic TEs also displayed markedly increased nascent transcription, further supporting that these regions represent active enhancers.

**Figure 3:**
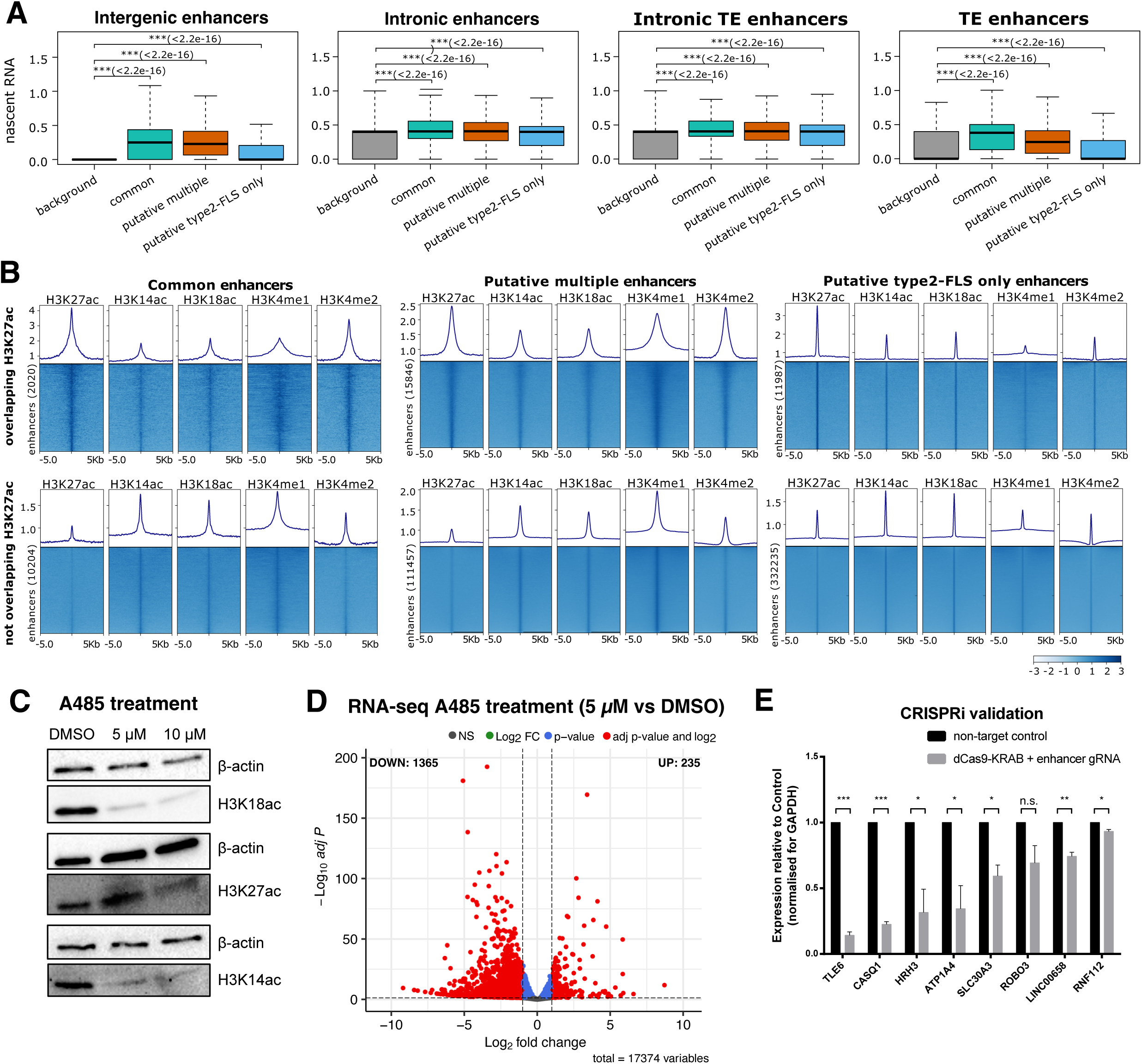
Validation of predicted enhancers. *(A)* Nascent transcription at different categories of enhancers: intergenic, intronic, intronic TEs and TEs. For each category of enhancers, we selected their corresponding background (see *Materials and Methods*). We also split enhancers into three categories: *(i)* common enhancers (detected by both type2-FLS and STARR-seq), *(ii)* putative multiple (detected by type2-FLS and other ML methods but not by STARR-seq) and *(iii)* putative type2-FLS only (detected only by type2-FLS). A Mann-Whitney U test was performed to evaluate whether the difference between the nascent transcription at the enhancer and corresponding background is statistically significant. Statistical comparisons within each panel were corrected for multiple hypotheses testing using the false discovery rate (FDR) method, and adjusted p-values are reported in the panels (p value: n.s. ≥ 0.05, * < 0.05, ** < 0.01 and *** < 0.001). *(B)* ChIP-seq signals for H3K18ac, H3K27ac, H3K14ac,H3K4me1 and H3K4me2 centred around enhancers. We considered separately the case of*: (i)* common enhancers, *(ii)* putative multiple and *(iii)* putative type2-FLS only. We also considered separately the case of enhancers overlapping H3K27ac peaks (top panels) or not (bottom panels).*(C)* Western Blot for H3K18ac, H3K27ac and H3K14ac upon A485 treatment (a P300 inhibitor). We considered DMSO and 2 concentrations for A485 (5 *µM* and 10 *µM*) and used β-actin as a control. *(D)* Volcano plot for changes in gene expression at 5 *µM* A485 compared to DMSO. *(E)* RT-qPCR result upon CRISPRi silencing of seven type2-FLS novel annotated enhancers. The experiment was performed in three technical replicates and three biological replicates, and a t-test was performed to evaluate whether the difference is statistically significant (p value: n.s. ≥ 0.05, * < 0.05, ** < 0.01 and *** < 0.001).

### Chromatin context of enhancers lacking H3K27ac

To further investigate whether these two classes of putative enhancers in H9 cells (using the H9 31 features trained models) display different chromatin contexts, we considered five histone modifications (H3K27ac, H3K14ac, H3K18ac, H3K4me1 and H3K4me2) at enhancers overlapping with H3K27ac peaks or not. Figure 3B shows that common, putative multiple and putative type2-FLS only enhancers that do not overlap H3K27ac display elevated H3K18ac and H3K14ac signals, suggesting that these acetylation marks may compensate for the absence of H3K27ac and contribute to enhancer activation. Additionally, putative type2-FLS only enhancers displayed stronger H3K4me2 enrichment relative to H3K4me1, consistent with H3K4me2 role as an enhancer-associated mark ^65^, which is in contrast to common and putative multiple enhancers.

### Strong depletion of H3K18ac leads to downregulation of gene expression independent of H3K27ac

Figure 3B confirms that predicted enhancers in H9 cells (using the H9 31 features trained models) display enrichment in H3K18ac, H3K27ac and H3K14ac. This raises the question of whether one of these acetylation marks is important or all of them are required for enhancers to be active. To test the importance of H3K18ac in enhancer regulation, we treated H9 hESCs with A485, a potent inhibitor of the histone acetyltransferase p300, and this treatment did not affect cell viability (Figure S7A). 24 hours of treatment with A485 leads to a substantial decrease (80%) of H3K18ac at 5 *µM*, without noticeable changes in the global levels of H3K27ac, which is only affected at 10 *µM* (Figure 3C and Figure S7B-D). It is worthwhile noting that there is also partial depletion of H3K14ac, albeit less than that of H3K18ac (approximately half). We then performed RNA-seq of 5 *µM* A485-treated and control H9 hESCs and found that 1,600 transcripts are differentially expressed, with most of them (1,365) being downregulated (Figure 3D). At 10 *µM*, we found only a small increase in the number of downregulated genes (Figure S7E), and most genes that are downregulated or upregulated at 5 *µM* of A485 are downregulated or upregulated at 10 *µM* of A485 (Figure S7F). Approximately 85% of the downregulated genes are proximal (within 25 Kb) to a predicted enhancer that the type2-FLS method labels as having medium or high levels of H3K18ac (Figure S7G), suggesting that those enhancers potentially control the proximal gene as previously reported ^66^. Our results show that the majority of downregulated genes are proximal to an enhancer that has both H3K18ac and H3K27ac, but the loss of H3K18ac at 5 *µM* of A485 is sufficient to downregulate the gene. In other words, the presence of H3K27ac at these enhancers is not sufficient to compensate for the loss of H3K18ac. Most importantly, one-third of the downregulated genes are proximal to a predicted enhancer that type2-FLS labels as having high levels of H3K18ac and low levels of H3K27ac (Figure S7H). This further supports that the presence of H3K18ac and not of H3K27ac is important for the enhancer activity at a subset of enhancers, and the loss of H3K18ac results in downregulation of proximal genes.

We further investigated the acetylation marks present at the promoters of these downregulated genes proximal to an H3K18ac-enriched enhancer and observed that all downregulated genes display high levels of H3K18ac and H3K14ac and no enrichment of H3K27ac (Figure S7I). This is in contrast to genes that are expressed and not differentially regulated, which also show enrichment of H3K27ac along with H3K18ac and H3K14ac (Figure S7I). Upregulated genes show similar patterns of enrichment as downregulated genes (Figure S7I), and one possibility is that their upregulation is an indirect effect of changes in expression levels of upstream regulators. Overall, our results show that downregulation upon A485 treatment is only observed at expressed genes that lack H3K27ac but have H3K18ac at both promoters and at a proximal enhancer.

### CRSIPRi targeting novel enhancers can lead to changes in expression of target genes

Since the downregulated genes have H3K18ac both at promoters and proximal enhancers (using the H9 31 features trained models), our A485 treatment results cannot confirm whether it is the loss of H3K18ac at enhancers or promoters that leads to their downregulation in H9 cells. To further investigate this, we have selected a set of 13 enhancers (corresponding to 13 proximal genes and 2 additional genes to which the enhancer is part of their introns, see Table S2) that we silenced using CRISPRi (dCas9 fused with a KRAB domain). Given that we target only the enhancer and not the promoter of the gene, this would allow us to determine whether the presence of H3K18ac at enhancers is functional or not. For each enhancer, we designed 3 sgRNAs and used a scrambled sgRNA as control. Our results showed that in the case of 6 enhancers (corresponding to 7 genes), we could see significant downregulation of the target genes by CRISPRi (see Figures 3E and S8A). In addition, silencing 7 enhancers leads to no change in gene expression (Figure S8B-C). We have two cases where enhancers were assigned to 2 genes. In the first case, silencing of an enhancer located within the intron of *ATP1A4* and near *CASQ1* genes leads to downregulation of both genes, indicating that the enhancer controls multiple genes. In contrast, silencing another enhancer located within the intron of the *LINC00658* gene leads to its downregulation, but the expression of the nearby gene, *PROKR2*, is not affected, indicating that in this case, the enhancer controls a single gene. Regulation of genes by enhancers is complex, with many scenarios where there are enhancer redundancies and, in some cases, complex regulatory logic ^67^, but we cannot exclude the case that some of the annotated enhancers are false positives.

### Seven epigenetic features are sufficient to predict enhancers in different human cell lines accurately

The application of any ML and AI models with 31 epigenetic features would be prohibitive due to costs, but also in systems where there is limited material available to perform the experiments. Hence, we took an unbiased approach and performed feature reduction to identify the core features that allow prediction of enhancers in H9 cells without a significant reduction in the performance of the ML/AI algorithms (see *Materials and Methods*). Our approach identified H2AFZ, H3K14ac, H3K18ac, H3K4me1, H3K4me2, H3K4me3 and ATAC-seq as the main features for human enhancers. We then trained ML/AI models in H9 cells using only these seven epigenetic features and observed that all except type2-FLS showed a small reduction in performance (AUCROC reduces by 0.02 or 0.03); Figure 4A. In contrast, type2-FLS displayed the same performance with the seven epigenetic features as with all 31 features, further supporting the robustness of the model. Most of the enhancers annotated with the 31 epigenetic features models are also annotated with the 7 features models (between 60%-90%); Figure S9A. All four models trained on 7 epigenetic features displayed a slight improvement in accuracy and recall (Figure S9B).

**Figure 4:**
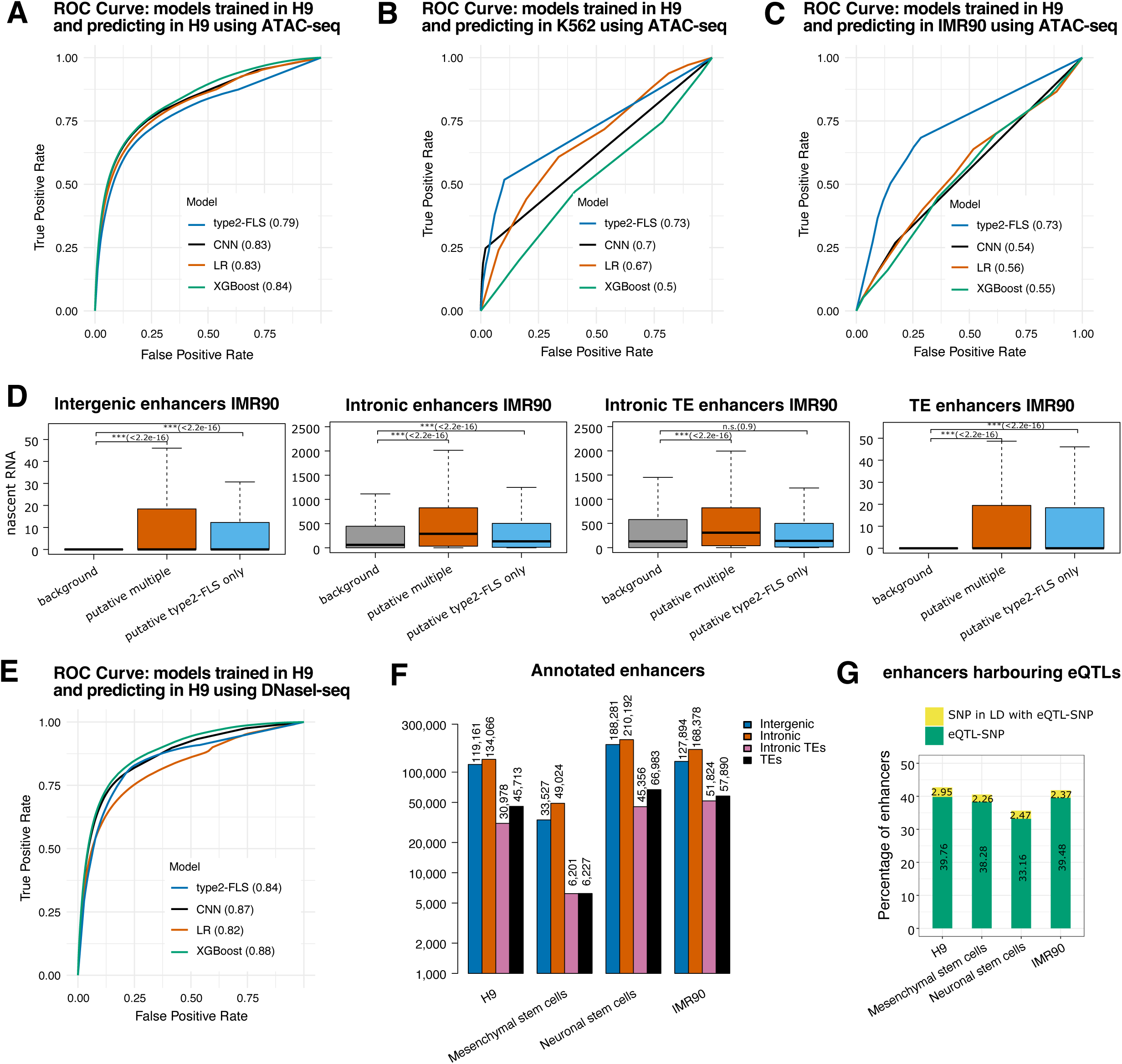
Minimal human model for enhancer annotation. *(A)* ROC curve and AUC values for four ML/AI models (CNN, LR, XGBoost and type2-FLS) trained on H9 human ES cells only on seven epigenetic features (H2AFZ, H3K14ac, H3K18ac, H3K4me1, H3K4me2, H3K4me3 and ATAC-seq). Training was performed on 500K bins, and the evaluation was performed on all unseen data. *(B-C)* Evaluation of ROC curve and AUC for all four models in two other cell lines *(B)* K562 and *(C)* IMR90 using Enhancer Atlas 2.0 annotated enhancers on the unseen data. *(D)* Nascent transcription at different categories of enhancers in IMR90 cells: intergenic, intronic, intronic TEs and TEs. For each category of enhancers, we selected their corresponding background (see *Materials and Methods*). We also split enhancers into two categories: *(i)* putative multiple (detected by type2-FLS and other ML methods) and *(ii)* putative type2-FLS only (detected only by type2-FLS). A Mann-Whitney U test was performed to evaluate whether the difference between the nascent transcription at the enhancer and the corresponding background is statistically significant. Statistical comparisons within each panel were corrected for multiple hypotheses testing using the false discovery rate (FDR) method, and adjusted p-values are reported in the panels (p value: n.s. ≥ 0.05, * < 0.05, ** < 0.01 and *** < 0.001). *(E)* Same as *(A)* but using DNaseI-seq instead of ATAC-seq. *(F)* Barplot with number of enhancers in four cell lines (H9, Mesenchymal stem cells, Neuronal stem cells and IMR90) using the type2-FLS enhancer annotation model with seven epigenetic features. We used the type2-FLS model with DNaseI-seq and split the enhancers into intergenic, intronic, intronic TEs and TEs. *(G)* The percentage of annotated enhancers that harbour at least one eQTL from GTEx database for each of the four cell lines. Green bars represent the percentage of enhancers that harbour at least one eQTL from GTEx database, while yellow bars represent the percentage of enhancers harbouring SNPs in LD with at least one eQTLs from GTEx database.

To evaluate if these models generalise well in other cells, tissues or disease conditions, (which were not used for training) we considered the enhancer annotation in K562 and IMR90 from Enhancer Atlas 2.0 ^60^ together with epigenetic profiles for the minimal model. Our result confirmed that type2-FLS performs consistently well in both K562 and IMR90 (AUCROC = 0.73 and AUCPR = 0.005 in K562; AUCROC = 0.73 and AUCPR = 0.374 in IMR90), further supporting the generalisation of this model (Figure 4B-C). CNN and LR also perform well, but only in K562 cells (AUCROC =0.7 and AUCPR = 0.141 for CNN and AUCROC=0.67 and AUCPR = 0.168 for LR), suggesting that they might generalise in some cells or tissues, but not in all. It is worthwhile noting the reduction in the performance of the models, even for type2-FLS (AUCROC reduced from 0.79 to 0.73). Differences in the relative contribution of specific epigenetic features may also partly reflect variability in ChIP-seq data quality rather than true biological divergence. Factors such as sequencing depth, mapping efficiency, signal-to-noise ratio, and replicate reproducibility can influence signal quantification, thereby affecting downstream model training and feature attribution.

One explanation for the reduction in model performance between cell lines is that the models can predict accurately housekeeping enhancers (predicted in multiple cells) in all cell lines, while it can predict cell type specific enhancers only in the cells the model was trained on (i.e., they cannot capture the cell type specific enhancers in other cells). To investigate if this is the case, we generated the ROC curve for the cell type specific enhancers in the two cell lines that the model was not trained on and observed only a small decrease in the AUC in K562 for type 2-FLS (AUCROC 0.7 for cell type specific enhancers) and no difference for the CNN model; see Figure S9C. In IMR90 there was a larger decrease in performance for cell type specific enhancers (AUCROC=0.58 for type2-FLS); see Figure S9D. This suggests that the model was not trained to predict only housekeeping enhancers and can predict cell type specific enhancers in most cell lines. This is not surprising since we have shown previously that there are negligible differences between the epigenetic code of housekeeping and developmental enhancers ^22^.

To further evaluate the accuracy of the predictions of our models, we split the predicted IMR90 enhancers into two groups: *(i)* putative type2-FLS only (enhancers only predicted by type2-FLS) and *(ii)* putative multiple (enhancers predicted by type2-FLS and other ML methods); see Figure S9E. Figure 4D shows that both groups of predicted enhancers display a higher level of nascent RNA compared to the corresponding genomic background, but the enhancers predicted by multiple methods display higher levels of nascent RNA. This indicates that all methods can detect the strong enhancers, but type2-FLS can also detect weak enhancers. That shows the improved generalisability of type2-FLS likely arises from its explicit modelling of uncertainty, which enables the system to capture weak enhancer signals that may be overlooked by more rigid models. As a result, the type-2 FLS provides a robust framework for detecting subtle regulatory patterns across diverse biological contexts.

### Characterisation of enhancers in multiple human cell lines

In some cell lines, ATAC-seq is not available, but DNaseI-seq, another method to measure DNA accessibility, was available, so we also trained four models with DNaseI-seq instead of ATAC-seq. This resulted in a slight improvement for the AUC of all methods except Logistic Regression, with the other models showing a higher AUC compared to even the 31 features model (Figure 4E). Overall, we found the type2-FLS model with 7 features, including DNaseI-seq performed well; Figure S9F. To keep consistency and due to data availability, we applied the 7-feature type2-FLS model with DNaseI-seq that was trained in H9 cells to predict enhancers in: H9, Neuronal Stem Cells, Mesenchymal Stem Cells and IMR90; Figure 4F. Our results show a similar distribution of enhancers into intronic, intergenic, and TE-derived enhancers. Furthermore, approximately half of H9 predicted enhancers are also found in Neuronal Stem Cells, and half of Mesenchymal Stem Cells predicted enhancers are found in Neuronal Stem Cells, while almost 70% of Mesenchymal Stem Cells predicted enhancers are found in IMR90 (Figure S9G). This is not surprising given that Mesenchymal Stem Cells and fibroblasts (including IMR90) are phenotypically similar ^68^. Most importantly, between 33% and 40% of these enhancers harbour an eQTL from GTEx ^69^ and, in addition, approximately 2.5% harbour a variant in strong LD with an eQTL, which provides additional evidence that those regions are enhancers, as assessed by changes in gene expression upon genetic variation within the enhancer sequence Figure 4G.

We then split enhancers into cell-type-specific (if they are predicted only in one cell type) and ubiquitous (if they are predicted in all five cell lines), with approximately 33K enhancers found in all cell lines used in our analysis; see Figure S9G. Our results showed that approximately one-fifth of enhancers overlap TEs, indicating that TEs might contribute to cell-specific gene regulation ^61^. To further investigate this, we considered cell-specific enhancers and ubiquitous enhancers and analysed whether any class of TE preferentially harbours these enhancers. Interestingly, Line 1 elements showed the highest diversity to cell-specific enhancers, with L1PA16 harbouring IMR90-specific enhancers, L1MB7 MSC-specific enhancers, L1ME1 at NSC-specific enhancers and L1M1 at H9-specific enhancers, while L1PA7 harbours at ubiquitous enhancers (Figure S9H). Furthermore, LTR12C, LTR13 and LTR33 harbour most cell-type-specific enhancers in IMR90, MSC, NSC and H9 cells. It is worthwhile noting that H9 STARR-seq enhancers where harboured predominantly by different TEs families compared to all predicted H9 enhancers (L1PA2, L1PA3, LTR12C and LTR7) (Figure S9H), suggesting these putative enhancers can propose novel TE families that could act as enhancers. In contrast, AluJb is the Alu element harbouring both the most cell-type-specific in each cell considered in our analysis and the most ubiquitous enhancers. Overall, our results support a model where the evolution of Line 1 and LTR elements could have contributed to cell-specific gene regulatory programs, while the evolution of the Alu element might have contributed to a lesser extent.

### Identification of enhancers in mouse ES cells

While our human model is generalisable in different cell types, we wanted to know whether we can derive similar models in other organisms and how those would compare to the human-specific model. To address this, we trained four models (type2-FLS, CNN, LR and XGBoost) on 17 mouse ENCODE epigenetic features and STARR-seq enhancer annotation in mouse ES cells (E14); see Figure 5A. Similarly, as in the case of the human cell lines, all of the models performed well (AUCROC between 0.81 and 0.87 and AUCPR between 0.1 and 0.204). Furthermore, the novel putative enhancers showed enrichment in similar epigenetic features to the ones detected by STARR-seq (Figure 5B). When investigating the rules for the type2-FLS, we found that H3K27ac is involved in more rules for enhancers in mouse ES cells, but H3K18ac is also involved in 10 of the top rules (Figure 5C and Figure S10). Interestingly, similar to human enhancer rules, the lack of DNA accessibility and methylated CpG is an indication of non-enhancers (Figure 5C and Figure S10). In addition, for mouse ES cells, low H3K27ac and H3K18ac also mark non-enhancer regions.

**Figure 5:**
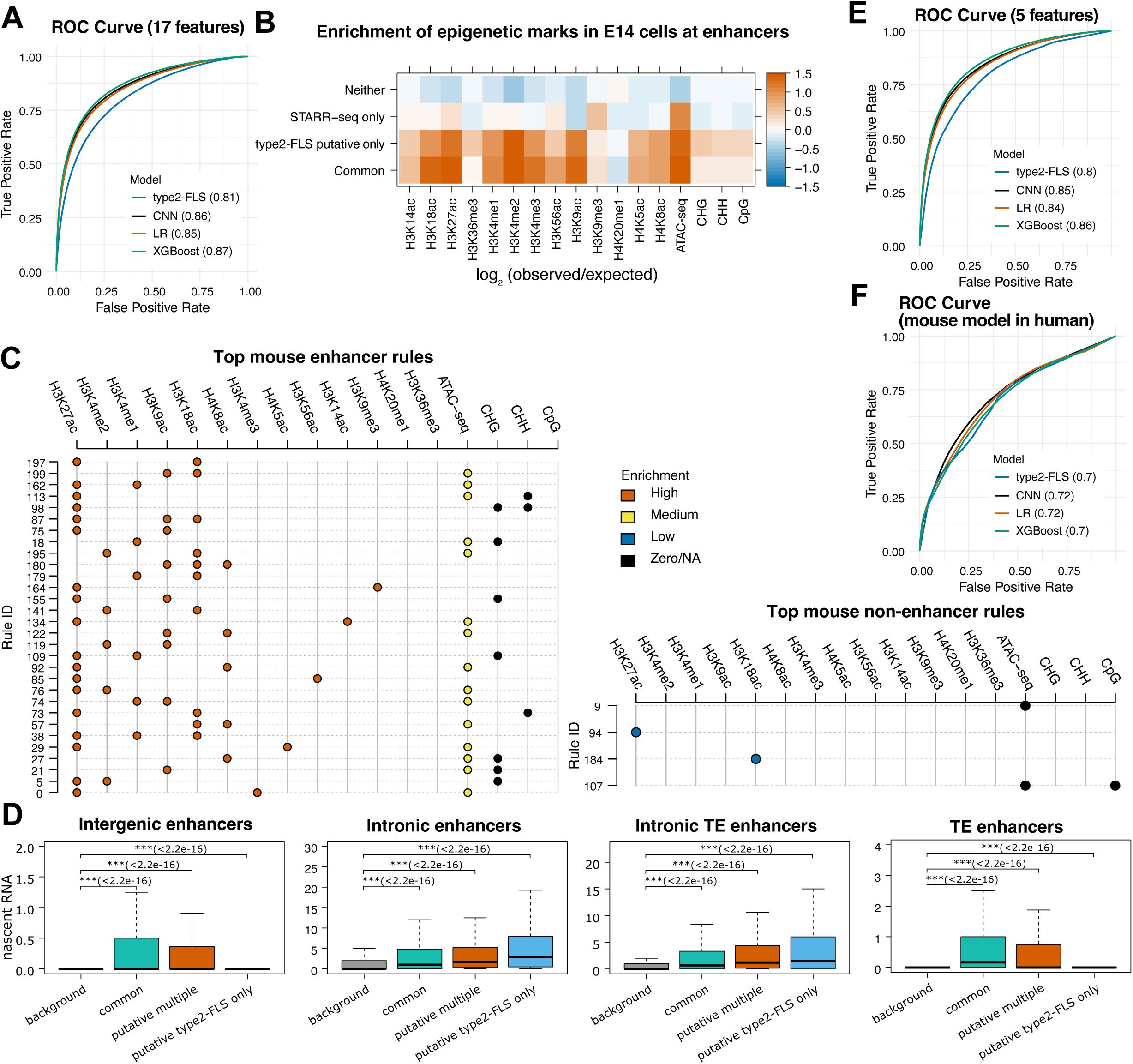
Mouse ML/AI models for enhancer annotation. *(A)* ROC curve and AUC values for four ML/AI models (CNN, LR, XGBoost and type2-FLS) trained on E14 mouse ES cells using 17 epigenetic features. Training was performed on 500K bins, and the evaluation was performed on all unseen data. *(B)* Enrichment of all 17 epigenetic signals (log_2_ observed/expected) at common enhancers (shared between type2-FLS and STARR-seq), putative enhancers (type2-FLS only), STARR-seq only enhancers and regions that are not classified as enhancers by either of the methods. *(C)* The top rules to classify enhancers (left) and non-enhancers (right). Each row represents a rule, and the colour represents the enrichment of the signal (red for high, yellow for medium, blue for low and black for zero/NA). *(D)* Nascent transcription at different categories of enhancers: intergenic, intronic, intronic TEs and TEs. For each category of enhancers, we selected their corresponding background (see *Materials and Methods*). We also split enhancers into three categories: *(i)* common enhancers (detected by both type2-FLS and STARR-seq), *(ii)* putative multiple (detected by type2-FLS and other ML methods but not by STARR-seq) and *(iii)* putative type2-FLS only (detected only by type2-FLS). A Mann Whitney U test was performed to evaluate whether the difference between the nascent transcription at the enhancer and corresponding background is statistically significant. Statistical comparisons within each panel were corrected for multiple hypotheses testing using the false discovery rate (FDR) method, and adjusted p-values are reported in the panels (p value: n.s. ≥ 0.05, * < 0.05, ** < 0.01 and *** < 0.001). *(E)* ROC curve and AUC values for four ML/AI models (CNN, LR, XGBoost and type2-FLS) trained on mESC using only five epigenetic features (H3K27ac, H3K4me1, H3K9me3, H3K18ac and ATAC-seq). Training was performed on 500K bins, and the evaluation was performed on all unseen data. *(F)* Evaluation of mouse minimal models in humans. ROC curve and AUC values of the four ML/AI models (CNN, LR, XGBoost and type2-FLS) trained on mES cells using only five epigenetic features to predict enhancers genome-wide in human ES cells on the unseen data.

Next, we merged individual bins into enhancers and selected the regions that are at least 200 bp and do not overlap with promoters. We recover the majority of STARR-seq annotated enhancers (∼20K) and detect approximately 374K additional enhancers, with only 9% previously reported in Enhancer Atlas 2.0 (Figure S11A). Similar to human ES cells, putative and common (with STARR-seq) enhancers are mainly intronic and intergenic, but approximately 20% are TE-derived enhancers (Figure S11B-D). Furthermore, we also found that many of the enhancers are within 1-10 Kb of each other (Figure S11E), with the total number of enhancers (402.7K) dropping to 242.6K when merging enhancers within 1 Kb and to 63.5K when merging enhancers within 10 Kb (Figure S11F).

Again, we split the predicted mESC enhancers into three groups: common enhancers (enhancers detected by both type2-FLS and STARR-seq), *(ii)* putative multiple (enhancers predicted by type2-FLS and other ML methods but not by STARR-seq) *and (iii)* putative type2-FLS only (enhancers only predicted by type2-FLS that do not overlap with the STARR-seq annotated enhancers); see Figure S11G. Figure 5D shows that both common enhancers and putative multiple display higher levels of nascent RNAs independent of whether the enhancer is intergenic, intronic, intronic TE or TE. Nevertheless, for intronic and intronic TE enhancers, putative type2-FLS only enhancers display the highest level of nascent RNA, indicating strong activity.

Common, putative multiple and putative type2-FLS only enhancers that do not overlap H3K27ac display elevated H3K18ac signals, suggesting that this modification could compensate for the absence of H3K27ac and contribute to enhancer activation (Figure S11I). While, in human H9 cells, we found H3K4me2 modification also located at putative type2-FLS only enhancers, in mESc, we only find this modification at putative multiple enhancers.

### Five epigenetic features are sufficient to accurately predict enhancers in multiple cell types or organisms

Finally, we investigated whether we could train ML/AI models in mouse ES cells with a smaller number of features to predict enhancers in other cell types or organisms. Our feature selection step identified a model with only six features (H3K27ac, H3K4me1, H3K9me3, H3K18ac, H3K20me1 and ATAC-seq), and all ML/XAI models (CNN, LR, XGBoost and type2-FLS) displayed good performance with an AUCROC of 0.80 or above (Figure S12A-C). H3K20me1 was not available in human and, thus, we decided to retrain a model with only five epigenetic features (H3K27ac, H3K4me1, H3K9me3, H3K18ac and ATAC-seq) in mESC, and the drop in performance was only negligible, where only LR drops the AUCROC from 0.85 to 0.84 (Figure 5E and Figure S12C). We evaluated this minimal model in a pre-T cell line in mouse (which was not used during model training), and the type2-FLS, LR and XGBoost performed well (AUCROC between 0.76 and 0.78), while CNN performed slightly worse (AUCROC=0.7), when compared to already annotated enhancers by ENCODE ^1^ (Figure S12E-G). Interestingly, we considered 24 different types of T cells, and for 14 of those, the performance was similarly high (AUCROC 0.76-0.77), indicating that there is a common regulatory program between some of the different subtypes of cells (Figure S12G). We further investigated if the models could predict cell type specific enhancers in the pre-T cells when they were trained in mESC and found that in there is no reduction in the prediction accuracy of cell type specific enhancers (see Figure S12H). Overall, this supports that the models trained on mESC epigenetic data have high accuracy in predicting enhancers in other cells and this is also the case for cell specific enhancers.

To further evaluate the generalisation of this minimal mouse enhancer model, we tested its prediction power in human ES cells, and our results confirmed that all four models trained in mouse ES cells on the five epigenetic features can accurately predict enhancers in human cells (with an AUCROC of 0.72 for CNN, 0.72 for LR and 0.70 for XGBoost and type2-FLS); see Figure 5F and Figure S12I. Nevertheless, only the LR model trained in human ES cells was able to generalise to mouse ES cells (with AUCROC = 0.6); see Figure S12J-K. We further investigated if the models could predict species specific enhancers, i.e., hESC specific enhancers when they were trained in mESC and vice versa. Our results show that there is a reduction in the prediction accuracy of species specific enhancers, with AUCROC between 0.59 and 0.69 for the mESC trained model and AUCROC between 0.57 and 0.63 for the hESC trained model (see Figure S12L-M). It is worthwhile highlighting that LR and type2-FLS performed consistently better across species including at identifying organism specific enhancers (with AUCROC between 0.59 and 0.72 and, for LR mESC trained model, an AUCROC of 0.69 when predicting hESC specific enhancers).

## Discussion

The majority of disease-associated mutations (more than 80-90%) are located in non-coding parts of the genome ^70,71^, with enhancers often harbouring more of these disease-associated mutations than promoters in some cases ^72^. Here, we applied different ML/AI methods and annotated enhancers in different cell lines in human and mouse using only epigenetic datasets (histone post-translational modifications, DNA accessibility and DNA methylation). Our models show high accuracy (with AUC greater than 0.7 and often even greater than 0.8) and identify hundreds of thousands of novel enhancers (from 140K up to 770K) with the majority of them being identified in a single cell/tissue, thus, indicating that they are cell/tissue specific. This raises the question of how many of these are indeed capable of driving expression. A recent study tested 680K regions (including enhancers in several cell types) and found that more than 40% are capable of driving expression, indicating that the number of enhancers is higher than previously reported and likely in the hundreds of thousands ^73^. Another study reported that up to almost 1M regions could act as enhancers, and one quarter are TE derived, which is within the range of the percentage we identify with our analysis ^61^. Furthermore, in our validation experiments by both global epigenetic perturbations and directed enhancer epigenetic rewriting, we show that indeed many of these novel enhancers are responsible for the activity of genes.

In addition, up to 43% harbour eQTLs or a SNP in strong LD with an eQTL, which provides further evidence that genetic variation within the enhancer region leads to changes in gene expression. In hESC (H9) cells, approximately 127K of the putative enhancers are detected by multiple ML/AI models (type2-FLS and another method) and 344K by type2-FLS only, with similar numbers found in mESC. The putative enhancers detected by multiple methods display significantly higher nascent RNA levels than background, independent of genomic context (intergenic, intronic, or TEs), indicating that they are active. In contrast, putative enhancers detected only by type2-FLS show medium and heigh nascent RNA levels only when located in introns. This pattern suggests that type2-FLS preferentially captures intronic regions with intermediate and potentially context-dependent enhancer activity that are missed by other models. Unlike logistic regression, which assumes additive and typically monotonic effects of each epigenetic feature, and unlike post-hoc explanations of opaque models that attribute importance to features one at a time, type2-FLS represents enhancer logic directly as fuzzy IF-Then rules that describe combinations of epigenetic marks in human-readable form. As a result, type2-FLS can both discover additional candidate enhancers with moderate or heterogeneous signals and simultaneously make explicit which specific combinations of chromatin and epigenetic features lead to these predictions. Altogether, these show that at least a large subset of the novel enhancers we identify are likely to be functional enhancers.

Opaque box models (such as CNN and XGBoost) can be interpreted, for example, using SHapley Additive exPlanations approach ^38^, while Logistic Regression can provide insights into the importance of different features by allowing examination of the coefficients. However, neither of these methods will be able to explain the combinatorial epigenetic code, e.g., a region has to have both enrichment of H3K27ac and H3K4me1 and depletion of H3K27me3 at the same time. SHAP and related post-hoc methods assign contribution scores to individual features or simple interactions; however, they do not by themselves yield a compact set of rules that exactly specify the multi-feature patterns used by the model for classification. This highlights the strength of our type2-FLS approach, which not only gives accurate predictions but also exposes human-readable relationships, such as IF H3K18ac is *High* and CpG methylation is *Medium*, then this will lead to an enhancer class. Crucially, these rules are part of the model itself rather than an external approximation, so they state precisely what the model has learned about enhancer-associated chromatin states. This is not possible by opaque box predictions or interpretable methods based on such opaque model induction (such as those using Shapley techniques). Most importantly, our type2-FLS based XAI approach allowed us to decode the combinatorial epigenetic code of enhancers and identify histone modifications that have not been usually linked to enhancer activity.

Our results showed that H3K18ac and H3K14ac are two post-translational histone modifications that are predictive of enhancers and may compensate at enhancers lacking H3K27ac. In particular, we showed not only that they are identified by eXplainable AI models as important (including their retainment in the subset of features for the minimal models), but also by global depletion of H3K18ac and partially of H3K14ac, leading to downregulation of a large number of genes (approximately 1.3K); Figures 3 and S6. Further simultaneous depletion of H3K18ac and H3K27ac only leads to downregulation of 300 additional genes (Figures 3 and S7), supporting the previous findings that H3K27ac is dispensable for gene activation in ES cells ^17,18^. In addition, silencing of enhancers enriched only in H3K18ac by CRISPRi led to downregulation of target genes in the case of approximately half of the tested enhancers (Figure 3E). Recent work has provided some insights into potential H3K18ac and H3K14ac mechanistic roles at enhancers. They showed that these marks promote accessibility of methyl writers and readers to H3K4, indicating a mechanism of crosstalk between H3K18ac/H3K14ac and H3K4 methylation in transcriptional regulation ^27,74^. Interestingly, this was not the case for H3K27ac.

Nevertheless, we do find enrichment of H3K27ac at many of our enhancers, but our results support a model where H3K27ac is not sufficient to identify all enhancers and, at the same time, not specific enough, so that all regions enriched in H3K27ac are annotated as enhancers. This was further supported by the rules identified by type2-FLS, where H3K27ac alone was not found in any of the top-identified rules in the human models. However, it is important to note that CRISPRi can induce silencing thousands of base pairs away from the guide RNA (gRNA) target site, which suggests that the regulatory effects observed via CRISPRi may not always be restricted to the specific local enhancer being targeted ^75^.

Furthermore, using a large repertoire of epigenetic modifications has been useful to identify epigenetic rules for enhancers, but using 31 epigenetic features is not practical to apply to many cells, conditions or disease states. Thus, by using feature selection, we identify a minimal set of seven epigenetic modifications that are sufficient to accurately predict enhancers in human cells and five epigenetic modifications for mouse cells. This opens new possibilities to further apply these models in multiple cell types, tissues, developmental stages and conditions using only a few ChIP-seq/CUT&Tag/CUT&RUN and ATAC-seq/DNaseI-seq datasets.

Most importantly, we showed that all four of the minimal models we trained in mouse ES cells (CNN, type2-FLS, LR and XGBoost) can generalise well in other mouse cells (pre-T cell line), but also in other organisms (human ES cells), that were not seen during training. Despite the lower AUCROC, type2-FLS demonstrated better generalisation across different cell lines. This suggests that while its absolute performance is reduced, it is capturing more stable and biologically transferable signals. The trade-off between peak predictive performance and cross-context generalisability highlights an important consideration for model selection, particularly in applications where robustness across biological systems is prioritised over maximum accuracy in a single dataset. Furthermore, while the type2-FLS model trained in human ES cells generalised well in multiple cell types not seen during training (and CNN and LR partially), LR was the only model trained in human ES cells that generalised to mouse ES cells. One possibility is that the generalisation failed for some models due to data quality and the pre-processing steps, but if this would be the case, generalisation should fail for all models, which is not the case. This indicates that the human-trained type2-FLS and CNN models can be applied to other cell types in humans, while all four models trained in mouse ES cells are more appropriate to be applied in multiple organisms. It is also worthwhile noting that the human model is trained on enhancer annotation in H9 cells from ChIP-STARR-seq, while the mESC model was trained on genome-wide STARR-seq. ChIP-STARR-seq focussed on sequences most likely to be enhancers (ChIP of several histone modifications associated with enhancers and several TFs specific to hESC), but it might miss potential enhancers enriched in other histone modifications ^12,13,76^. In contrast, the mESC STARR-seq has less bias given that it is genome-wide, and this can explain why it generalises better.

Finally, while type2-FLS may not always outperform other methods, it remains useful method for identification of enhancer candidates across varying activity levels, generalises relatively well in most other cell lines unseen during training compared to other ML methods and highlights the relevance of H3K18ac in enhancer biology. Overall, combining multiple ML models (including type2-FLS) would provide more flexibility in identifying candidate enhancers. In particular, selecting regions predicted to be enhancers by at least one of these models provides a more comprehensive set of candidate enhancers, while selecting the common set of enhancers predicted by multiple methods could represent a more conservative set of the strongest candidate enhancers.

## Materials and Methods

### Cell culture and CRISPRi

The H9 hESC line was obtained from WiCell. hESCs were grown on vitronectin-coated plates (Thermofisher, A14700) in mTeSR Plus medium (Stem Cell Technologies, 100-0276) supplemented with 100 U ml–1 penicillin–streptomycin (Gibco, 15140122) and passaged every 3–4 days with 0.05 mM EDTA (Invitrogen, 15575020), according to the manufacturer’s protocols.

Cells used for protein extraction for Western Blotting were incubated for 24h in media supplemented with 0, 5, 10 or 20 µM of A485 (Bio-Techne, 6387) in DMSO before being retrieved.

Cells used for CRISPRi were grown in eTeSR medium (Stem Cell Technologies, 100-1215) for at least one passage before nucleofection. Nucleofection was performed in an Amaxa 4D-Nucleofector (Lonza, V4XP-3032) according to the manufacturer’s protocol for the Human Stem Cell Line H9. We have generated sgRNA for enhancers using the offTargetAnalysis function of CRISPRseek ^77^ and selected the top 3 sgRNAs that did not have off-targets (see Supplementary Table S2). The sgRNAs were cloned on the PB_rtTA_BsmBI plasmid (Addgene ID 126028) ^78^ using BsmBI. For each enhancer region, the sgRNAs plasmid mixes (of equal amounts of each of the 3 sgRNAs targeting an enhancer), the CRISPRi-Bac plasmid (PB_tre_dCas9_KRAB, Addgene ID 126030) ^78^, and the piggyBac-transposase plasmid ^78^ were mixed in a 1:1:2 ratio for a total of 1 μg, and 2 x 10^5^ cells were nucleofected. A media exchange was performed 24 hours after nucleofection to remove ROCKi, and 48h following nucleofection, cell selection was started by supplementing the media with 40 μg ml–1 hygromycin B and 40 μg ml–1 of G418 for 10 days (media changed every other day). After selection and expansion, the CRISPRi system was activated by media supplementation with 500 ng ml–1 of doxycycline (Dox). Cells were collected for RNA isolation and RT-qPCR 48 h after Dox induction.

### Cell viability assay

Cell viability was measured using the MTS Assay kit (Abcam, #197010), according to the manufacturer’s instructions. 24 hours after treatment with A485 (concentrations ranging 0.2-50µM), the MTS reagent was added to the cell culture media and incubated for 1.5h under standard culture conditions. Absorbance was measured at 490nm after brief mixing.

### Western Blot

Cells were pelleted by centrifugation at 300g for 3 min and resuspended in cold RIPA buffer (150 mM sodium chloride, 1.0% Nonidet P40, 0.5% sodium deoxycholate, 0.1% SDS (sodium dodecyl sulfate) and 50 mM Tris, pH 7.4), supplemented with 1X Roche cOmplete EDTA-free protease inhibitor, and incubated for 30 min on ice with intermittent mixing. Samples were quantified using a Pierce™ BCA Protein Assay Kit (ThermoFisher, 23227).

Equal amounts of protein extract were denatured in 4x Laemmli Sample Buffer (BioRad, 1610747) and separated on Mini-Protean TGX Gels 4-20% (BioRad, 4561093), blotted on a nitrocellulose membrane (BioRad, 1620145) and immunoblotted with antibodies for H3K14ac (Abcam, ab52946, 1:500 dilution),

H3K18ac (Abcam, ab1191,1:1000 dilution), H3K27ac (Active Motif, 39133, 1:1000 dilution), β-actin (Abcam, ab8226, 1 µg ml–1 dilution) and H3 (Active Motif, 61799, 1:500 dilution), and horseradish peroxidase (HRP)-conjugated goat anti-rabbit IgG H&L (Abcam, ab6721) and HRP-conjugated goat anti-mouse IgG (BD Pharmingen, 554002) secondary antibodies.

### RNA-seq and RT-qPCR

Total RNA was isolated from H9 hESCs using TRIzol reagent (ThermoFisher Scientific, 15596026), according to the manufacturer’s instructions. The RNA was subsequently treated with DNaseI (NEB, M0303) and ethanol precipitated. RNA was assessed qualitatively and quantitatively using Quibit and Bioanalyzer 2100 (Agilent). For RNA-seq, poly(A) RNA selection, library preparation, and sequencing were carried out in three biological replicates by Novogene.

For RT-qPCR, the cDNA was prepared using RevertAid Reverse Transcriptase (ThermoFisher Scientific, EP0441) and Oligo(dT)18 (ThermoFisher Scientific, SO131) as primers. qPCR was performed using KAPA SYBR FAST qPCR Master Mix (2X) ABI Prism (Kapa Biosystems, KK4604) in a StepOnePlus Real-Time PCR System (Applied Biosystems). The list of specific primers used is given in Supplementary Table S3. RT–qPCR was done in triplicate, with three independent biological replicates and normalised to GAPDH.

### RNA-seq preprocessing

The adapters and low-quality reads have been trimmed using Trimmomatic ^79^, and Tophat2 ^80^ was used to align the reads to the hg38 genome reference. We used Picard tools ^81^ to deduplicate reads, HTseq ^82^ to count reads, and then DESeq2 ^83^ to detect differentially expressed genes. For DESeq2, we selected transcripts with at least 10 reads and used a P-value threshold of 0.05 and a log2FC threshold of 1.0. Preprocessing statistics for the RNA-seq data can be found in Supplementary Table S4. Supplementary Table S5 contains the results of DESeq2 analysis for A485 5 µM compared to DMSO treatment and Supplementary Table S6 contains the results of DESeq2 analysis for A485 10 µM compared to DMSO treatment.

### Human datasets retrieval and pre-processing

The Homo Sapiens genome (hg38) ^84^ was tilted into 100bp bins with the removal of sex chromosomes to avoid the potential biases arising from X inactivation mechanisms. A pre-processed (fold change over control) ChIP-seq, ATAC-seq, DNaseI-seq and Bisulfite data (CpG, CHH and CHG) dataset of 31 histone marks was downloaded from ENCODE for H9 ^85^; see Supplementary Table S7. For cell lines: IMR90 cells, mesenchymal stem cells (MSC), K562, and neuronal stem cells (NSC), 7 epigenetic marks were downloaded from ENCODE, which include H2AFZ, H3K14ac, H3K18ac, H3K4me1, H3K4me2, H3K4me3 and ATAC-seq/ DNaseI-seq. Both ATAC-seq and DNaseI-seq were available for H9, IMR90 and K562 cells, while for the other 2 cell lines (MSC & NSC) only DNaseI-seq was available. The enrichment score of each histone mark was normalised using min-max (values between 0 to 1) normalisation methods to investigate the most optimal and stable model performance on the test dataset; see Supplementary Figure S13A. Note that the optimal normalisation method was selected to ensure the lowest difference between recall of enhancers and non-enhancers with highest average recall. It is important to acknowledge that the use of the test data to select this normalisation method introduces a potential risk of overfitting to the test dataset. Nevertheless, the fact that the models can generalise in other cell lines and that the performance is reported on all unseen data (all data that was not used for training or testing) indicates that they were not overfitted. Values of 0 were replaced with NAs. Self-transcribing active regulatory region sequencing (STARR-seq) data for H9 were obtained from ^45^.

### Mouse datasets retrieval and pre-processing

The 17 histone modification datasets of all ChIP-seq, DNaseI-seq, ATAC-seq and WGBS sequencing were retrieved from GEO; see Supplementary Table S8. Firstly, *Trimmomatic* ^79^ was used to filter out the short and low-quality reads. Next, *Bowtie2* ^86^ was used to align the samples against their corresponding reference genome. It is worthwhile noting that inclusion of multi-mapping reads could lead to false positive peaks in TEs (affecting both model training and evaluation for TE enhancer prediction), while their exclusion could lead to false negative peaks in TEs. Here, we selected to proceed with inclusion of multi-mapping reads. *Macs2* ^87^ *narrow peak* software was used to generate the peaks for each sample. For ChIP-seq, to compute the fold change of IP over control, *deep tools bamCompare* ^88^ package was used with a binSize 20 and –operation parameter equal to ratio. For ATAC-seq and DNaseI-seq, *deeptools bamCoverage* ^88^ package was used with a binSize 20 and –operation equal to the ratio parameters. For whole genome bisulfite sequencing data pre-processing *bismark* ^89^ The tool was used to get the methylation proportion at each CpG, CHH and CHG site in the genome. For datasets in mm9, we used *liftover* to obtain mm10 coordinates.

After generating the bigwig files of all the samples, the 100bp Mus musculus genome (mm10) tiled was created, and we followed the same steps as for human analysis. For ChIP-seq samples, the tracks having a score of zero were changed to NA, and then all data were normalised by using the quantile normalisation (0.01 for min and 0.99 for max), which ensures the lowest difference between recall of enhancers and non-enhancers and the highest average recall on the test data; see Supplementary Figure S13B.

For the mouse pre-T cell line, five epigenetic features were selected (H3K18ac, H3K27ac, H3K4me1, H3K9me3, and ATAC-seq) were downloaded from NCBI see Supplementary Table S8, and for enhancer annotations, we have tried 24 different immune cell types from ENCODE and first selected the ones that showed the highest AUC for the ROC and, from those, the closest to pre-T cells, i.e., “naive thymus-derived CD4-positive, alpha-beta T cell”.

### Nascent transcription datasets and backgrounds

For nascent transcription levels, we used the following datasets: precision nuclear run-on sequencing (PRO-seq) in H9 hESC cells (GSE278359), bromouridine sequencing (Bru-seq) in IMR90 cells ^90^ and Transient Transcriptome sequencing (TT-Seq) in mESC cells^91^.

We computed the nascent transcription backgrounds for each category of enhancer: intergenic, intronic, intronic TEs and TEs. In particular, for intronic and intronic TEs enhancers, background transcription levels were computed at a similar number (to the number of intronic or intronic TEs putative enhancers) of 200bp bins that were not labelled as enhancers within introns of the corresponding genes harbouring STARR-seq enhancers. For intergenic enhancers, the background was computed at a similar number (to the number of intergenic putative enhancers) of random intergenic 200bp bins that do not overlap TEs, while for TEs enhancers, the background was computed using a similar number (to the number of TEs putative enhancers) of random 200bp bins that overlap TEs.

### Type-2 Fuzzy Logic System Framework

An interval type2 fuzzy logic system (type2-FLS) was implemented to identify and classify enhancers ^22,42^. Interval Type2-FLS employ interval type-2 fuzzy sets which can be represented in terms of upper and lower membership functions (forming a Footprint of Uncertainty - FOU), which enables the model to capture noise and uncertainty inherent in biological datasets. The type2-FLS mainly consists of five stages: *(i)* Fuzzification, transforming numerical inputs into interval membership values; *(ii)* Rule Base, defining fuzzy IF/THEN relationships between input and output variables; *(iii)* inference engine, computing rule firing strengths; *(iv)* Final Classification phase, yielding binary enhancer predictions (Enhancer/No Enhancer).

Input variables (e.g. epigenetic markers such as H3K4me1, H3K27ac, or H3K18ac) were modelled using interval type2 fuzzy sets with linguistic terms (zero/NA, low, medium and high). The antecedents of fuzzy rules take the form ‘’IF H3K4me1 is High AND H3K27ac is low and H3K18ac is medium” and the rule consequent is the the enhancer class (Enhancer/No Enhancer). An initial random rule base see Table S9 (maximum three antecedents and one consequent) was optimised using a multi objective multi constraint genetic algorithm (to maximise explainability via generating the smallest rules base of short IF-Then rules to generate the possible highest accuracy thus allowing to generate fully explainable models which are close in accuracy to black box models). The inference engine produced interval-valued outputs, from which support, confidence and dominance score (see equations 1-3) were computed to prioritise highly confident enhancer classification rules. The top rules are chosen based on the dominance value, where the higher the dominance, the higher the trust in the given rule

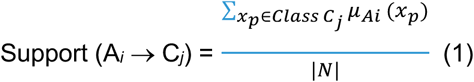

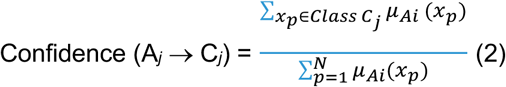

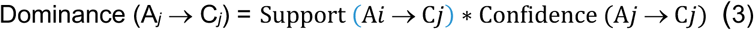

Where, *C_j_* is the *j*-th class in the set of predefined classes; μ_*Ai*_(*x_p_*) is the matching degree of each training example *x_p_* with the antecedents of rule *i*, where *p* = 1, …, *N*; *N*, the total number of samples in the dataset.^21,40^

### ML/AI models training and validation

For the model generation phase, we employed 500,000 bins in both the training and testing datasets, which were sampled from a pool of 3 million tiled bins (see Figure 1). We have also tested the overall generated model over the unseen out-of-sample remaining 29 million bins. The sample size (for training and testing data) maintains the proportional size of each chromosome labelled bin. To keep the same proportion of positive bins as for the genome-wide data, both training and testing datasets of human H9 cell line contain 496,189 randomly sampled non-consecutive bins as the (0) non-enhancer class and 3,813 randomly sampled non-consecutive bins as the (1) enhancer class. The training dataset was further divided into training and validation subsets in an 80:20 ratio. The validation set was utilised for hyperparameter tuning of the ML/AI models and early stopping to avoid overfitting the model. To handle the big imbalance between the enhancers/non-enhancers classes for the type-2 fuzzy XAI-based model, we have used the techniques reported in ^52^ to handle the class imbalance and generate highly accurate models with a small number of short IF-THEN Rules, which could be easily analysed, understood and augmented by the stakeholders. To handle the big imbalance between the enhancers/non-enhancers classes for the type-2 fuzzy XAI-based model, we have used the techniques reported in ^52^ to handle the class imbalance and generate highly accurate models with a small number of short IF-THEN Rules. For CNN, LR and XGBoost, the SMOTE class imbalance technique did not improve the performance, so we trained these models without any class imbalance technique. We implemented the type2 Fuzzy Logic Systems (type2-FLS), Convolution Neural Network (CNN), Logistic Regression (LR) and XGBoost models by using Python Libraries (i.e., Ex-Fuzzy ^92^, torch ^93^, scikit-learn ^94^ and XGboost ^95^). The hyperparameters for each model were optimised using GridSearchCV, which selected the optimal configuration based on cross-validation performance conducted on the training dataset; see Supplementary Table S10. Training was conducted using batch learning, with an early stopping criterion, for model regularisation during model training, implemented to enhance efficiency and prevent overfitting. A patience criterion of 3 was applied, halting the training process if no improvement was observed after three consecutive epochs, within a maximum of 20 epochs. All models were trained on the binary classification labels obtained from STARR-seq enhancer annotations, which we considered the ground truth. The best-fitted models were chosen based on three parameters, i.e., accuracy, average recall, precision and nonparametric ROC (Receiver Operating Characteristic) curve achieved on the training data sets. These models were then used to call the enhancers on all other cell lines. The predicted enhancers on the whole genome of the Human model were then filtered based on the prediction probability of 0.8 and higher (31 Feature model), 0.76 and higher (7 Feature model) and for the Mouse Model probability of 0.71 and higher (17 Features) was used. These thresholds were selected as the average between the optimal threshold for the nonparametric ROC curve (Youden’s J statistic) and the optimal threshold for the PR curve (F1 score); see Supplementary Table S11. These filtered regions were merged with their upstream and downstream region with a gap of 500bp; see Supplementary Figure S14. To annotate enhancers (e.g., intronic, intergenic or TE derived) we used the genomic annotation from UCSC Genome Browser ^96^.

All models were evaluated on the testing dataset (unseen 500,000 bins) and all unseen out-of-sample datasets (approximately 29 million bins for human and 24 million bins for mouse); Supplementary Table S1. In the manuscript, we report only the results for the unseen out-of-sample datasets, while results for the testing dataset can be found in Supplementary Table S1.

### Interpretation or explainability of ML/AI models

To interpret the Opaque box models (CNN and XGBoost) we used correlation Shapley Additive eXplanations (SHAP) ^38^. Specifically, for SHAP TreeExplainer (XGBoost), DeepExplainer (CNN) has been implemented in Python. The models were then analysed using these explainers to derive feature importance rankings. For Logistic Regression, we reported the correlation coefficients ^97^.

To ensure a high degree of predictive power along with explainability, for each type2-FLS model, a maximum of 100 rules were generated, and each rule contained up to three histone modifications. The epigenetic markers are individually divided into zero/NA, low, medium, and high classes without defining a stringent threshold, but allow the overlap between the classes to capture the uncertain regions of enhancers. These class boundaries vary by epigenetic mark and were computed while training the model.

### Feature selection method

Feature selection was conducted using the wrapper-based Recursive Feature Elimination (RFE) algorithm ^98^. Briefly, RFE iteratively trains a predictive model on the full feature set (“Base Features”), ranks variables according to their contribution to model performance, and removes the least informative feature(s) at each step. This recursive process continues until the predefined target number of features (set to 1) is reached. At every iteration, model performance is re-evaluated, producing a sequence of nested feature subsets and corresponding performance estimates, as well as feature importance scores reflecting relative predictive relevance. We selected seven features for human model and five features for mouse model to ensure that there is not a large drop in the corresponding performance estimate (less than 1% drop in recall).

### Enriched 3D contacts and Expression Data

Hi-C data in H9 cells from ^99^ was used to find the 3D contacts of potential enhancers. Promoter locations were defined as being regions up to 1 Kb upstream of TSS sites^94^. Enhancer bins that form a chromatin loop with the promoters located at more than 100 Kb were classified as distal, while enhancers located within 100 Kb of a promoter were labelled as proximal. For H9 cells, expression data were obtained from ^100^.

### Selection of H3K18ac-positive and H3K18ac-only enhancers

To characterise enhancers classes based on the epigenetic landscape, we defined membership thresholds derived directly from the optimised parameters of our type2-FLS. We categorised *loci* as H3K18ac-positive if they displayed a normalised signal for H3K18ac of at least 0.034 and as H3K18a only if they display a normalised signal for H3K18ac of at least0.034 and a normalised signal for H3K27ac of at most 0.01. This classification effectively separated enhancers enriched for H3K18ac and H3K27ac from those showing only enrichment for H3K18ac.

### eQTL analysis

To generate a comprehensive set of regulatory variants, all significant variant–gene pairs from the GTEx v8 eQTLs dataset were downloaded from the *GTEx* portal ^69^. To include variants in strong LD with eQTLs, we used the ensemblQueryR ^101^ package to query LD information through the Ensembl REST API. For each SNP, LD expansion was performed using the *encephalQueryLDwithSNPwindowDataframe* function, applying stringent LD thresholds (r² ≥ 1.0 and |D′| ≥ 0.8) within a ±500 bp window. LD information was obtained for the 1000 Genomes Project European population (EUR).

## Data and materials availability

All RNA-seq datasets from this study have been submitted to the NCBI Gene Expression Omnibus (GEO; http://www.ncbi.nlm.nih.gov/geo/) under accession number GSEXXXXXX. The scripts to perform the analysis and enhancer annotations for human and mouse cell lines can be accessed on Zenodo at https://doi.org/10.5281/zenodo.20326219 ^102^.

## Acknowledgments

We thank members of the Centre for Epigenetics at QMUL and Dr Alex Radzisheuskaya for discussion and feedback on this manuscript. The analysis was performed on the HPC at the University of Essex, and we thank Stuart Newman for his support in using the cluster.

## Author contributions

H.H. and N.R.Z. conceived and designed the analysis and experiments. K.M., J.W., C.G.C., R.A.B. and O.A.G. performed the computational work. D.P.B., B.K. and F.B. performed the experiments. N.R.Z., H.H., P.M., G.F. and C.G.B. supervised the work. K.M., D.P.B., H.H. and N.R.Z. wrote the paper. The authors read and approved the final manuscript.

