## Supplementary material for "Explainable AI identifies H3K18ac as a new marker of active enhancers": Figure S

[†]Co-first authors

[*]Corresponding authors

### Supplementary Figures

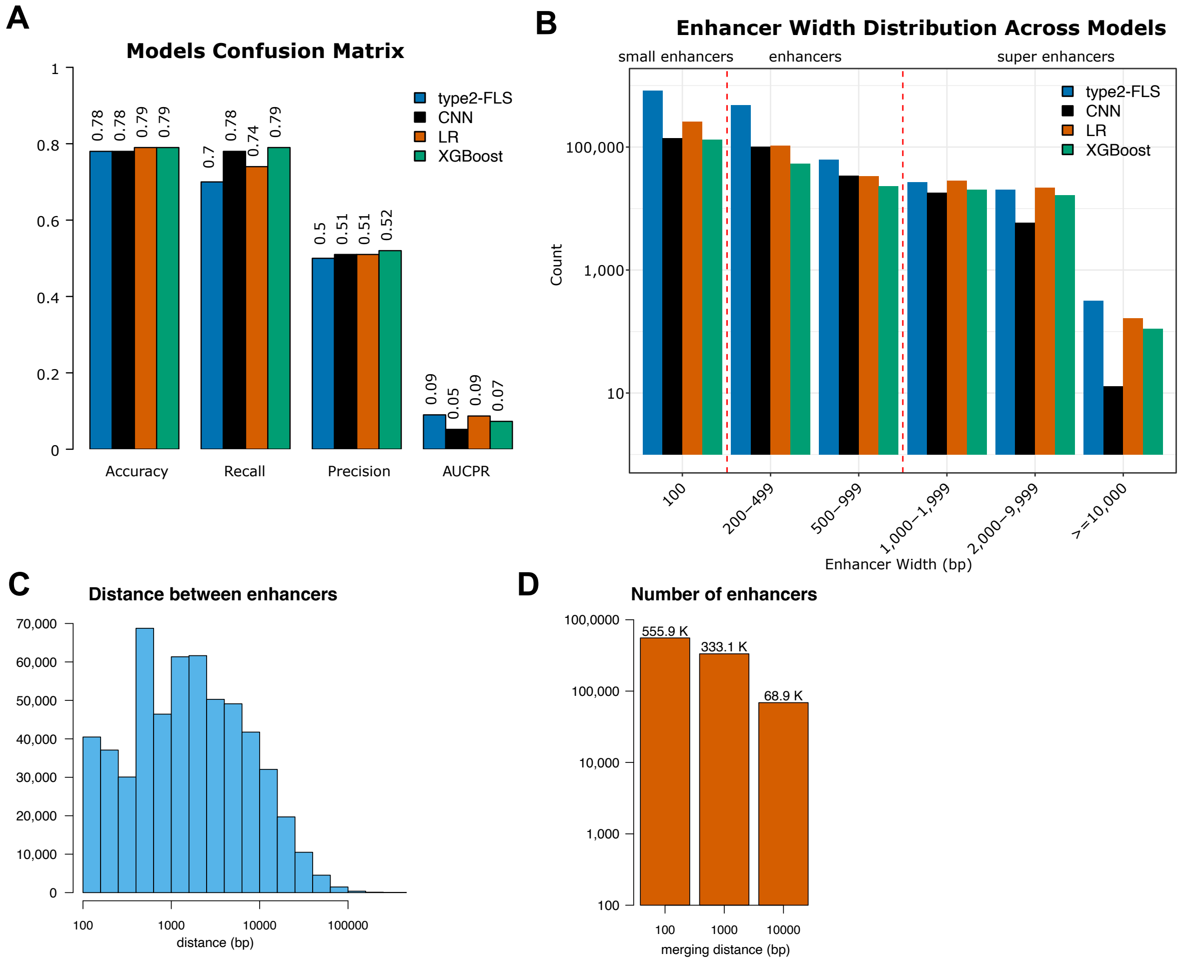


**Figure S1:** *Characterisation of enhancers predicted by the different ML/AI models in human ES cells. (A)* Confusion matrix for all four ML/AI models (CNN, LR, XGBoost and type2-FLS) trained in human ES cells. The values are reported on all unseen data. *(B)* Distribution of enhancer sizes for each of the four ML/AI methods. *(C)* Histogram of the distances to the nearest enhancer. *(D)* Number of enhancers after merging all enhancers within different distances of each other (100 bp, 1 Kb and 10 Kb).

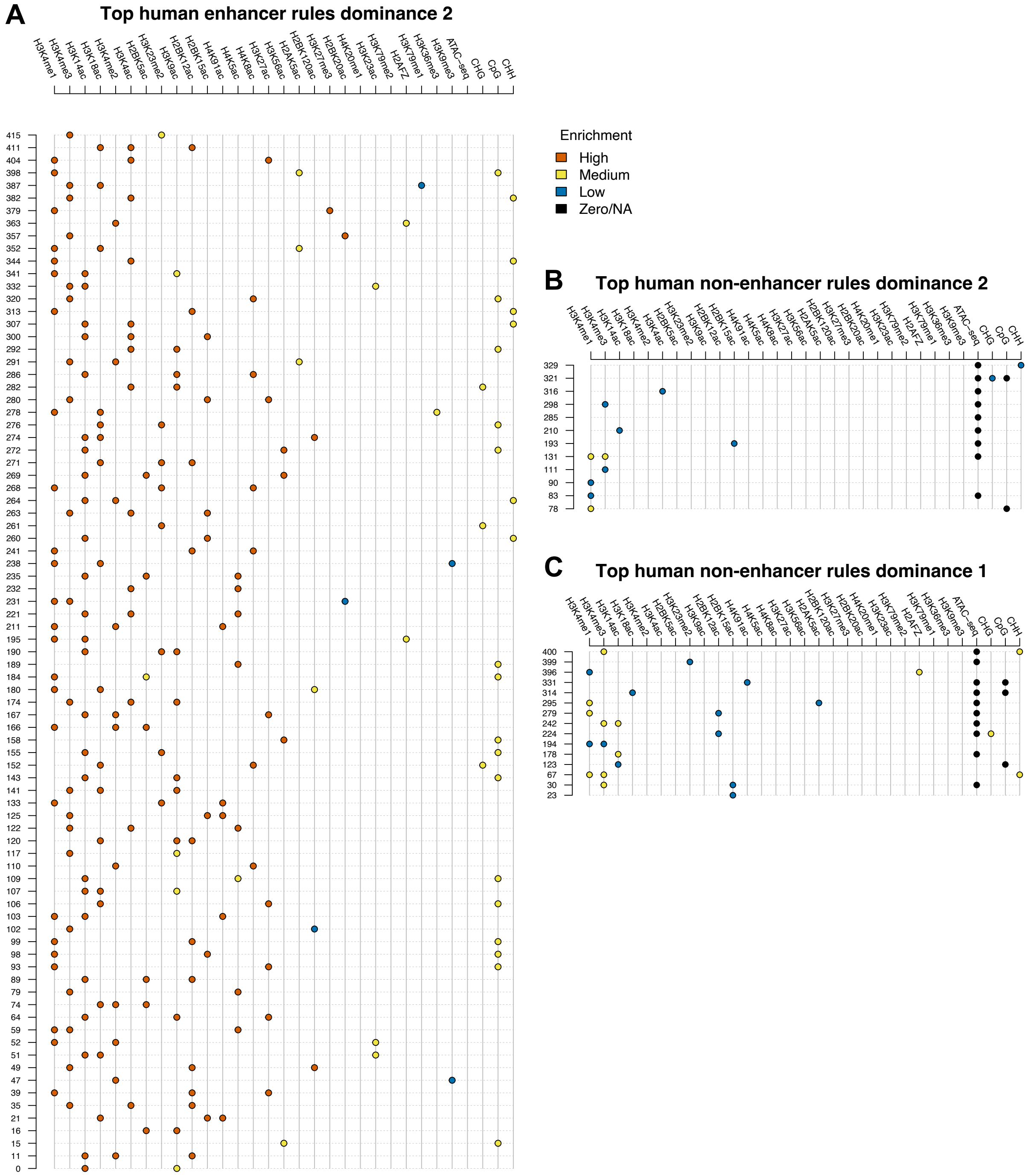


**Figure S2:** *Rules to annotate human enhancers.* We plot: *(A)* top human enhancer rules with dominance 2, *(B)* top human non-enhancer rules with dominance 2 and *(C)* top human non-enhancer rules with dominance 1. Each row represents a rule, and the colour represents the enrichment of the signal (red for high, yellow for medium, blue for low and black for zero/NA). For example, rule 352 is regions with a high level of H3K4me1, high levels of H3K18ac, and medium levels of H2AK5ac are annotated as enhancers.

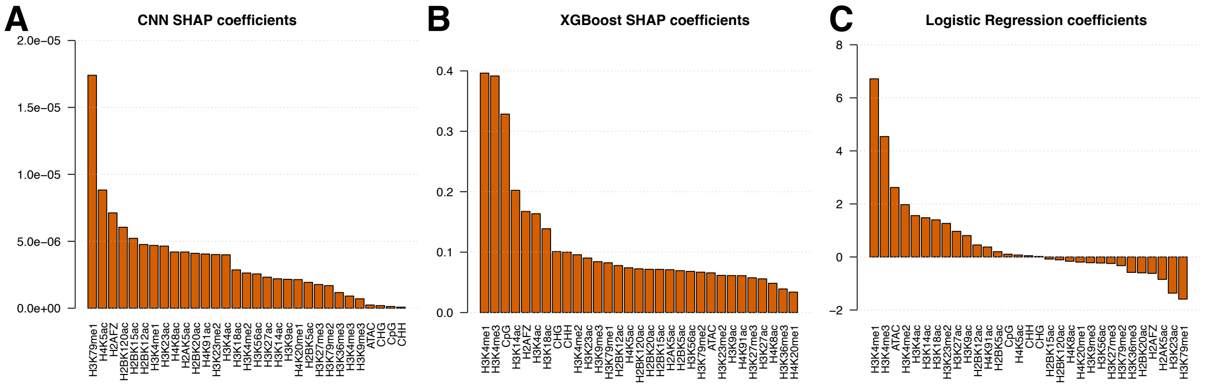


**Figure S3:** *Interpretation of ML/AI methods for the human ES cells. (A-B)* Barplots with the SHAP coefficients ordered decreasingly for *(A)* CNN and *(B)* XGBoost. Higher values indicate the corresponding feature is more important in the prediction of the corresponding model. *(C)* The coefficients of the Logistic Regression model. Strong positive and negative values indicate coefficients that are important for the predictions.

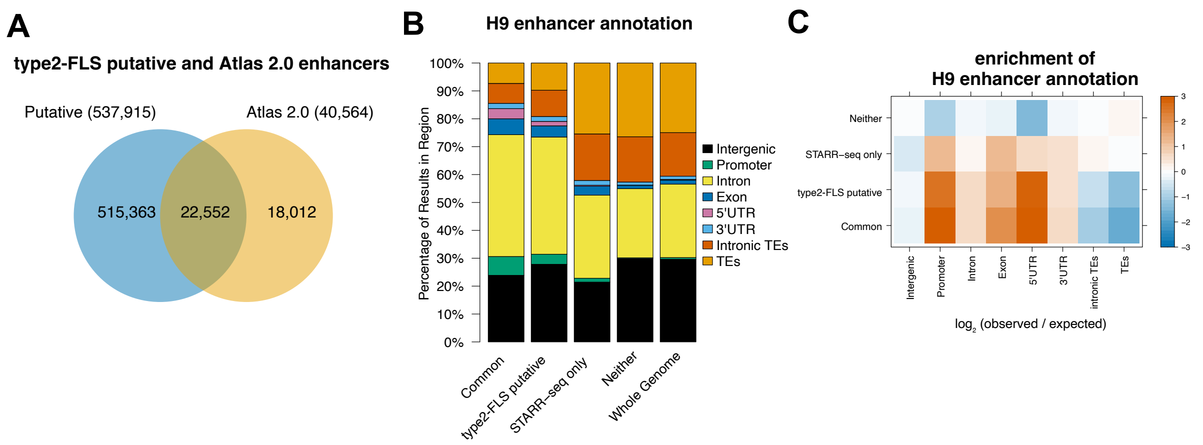


**Figure S4:** *Characterisation of human ES cells' putative enhancers. (A)* Overlap of putative enhancers with enhancers annotated in Enhancer Atlas 2.0. (*B)* Annotation of common enhancers (shared between type2-FLS and STARR-seq), putative enhancers (type2-FLS only), STARR-seq only enhancers and regions that are not classified as enhancers by either of the methods. We considered intergenic, promoter (up to 1Kb upstream of TSS), intron, exon, 5'UTR, 3'UTR, intronic TEs and TEs and plotted whole genome distribution of the features. *(C)* $\log_2$(observed/expected) overlaps based on the whole genome distribution of the different annotations.

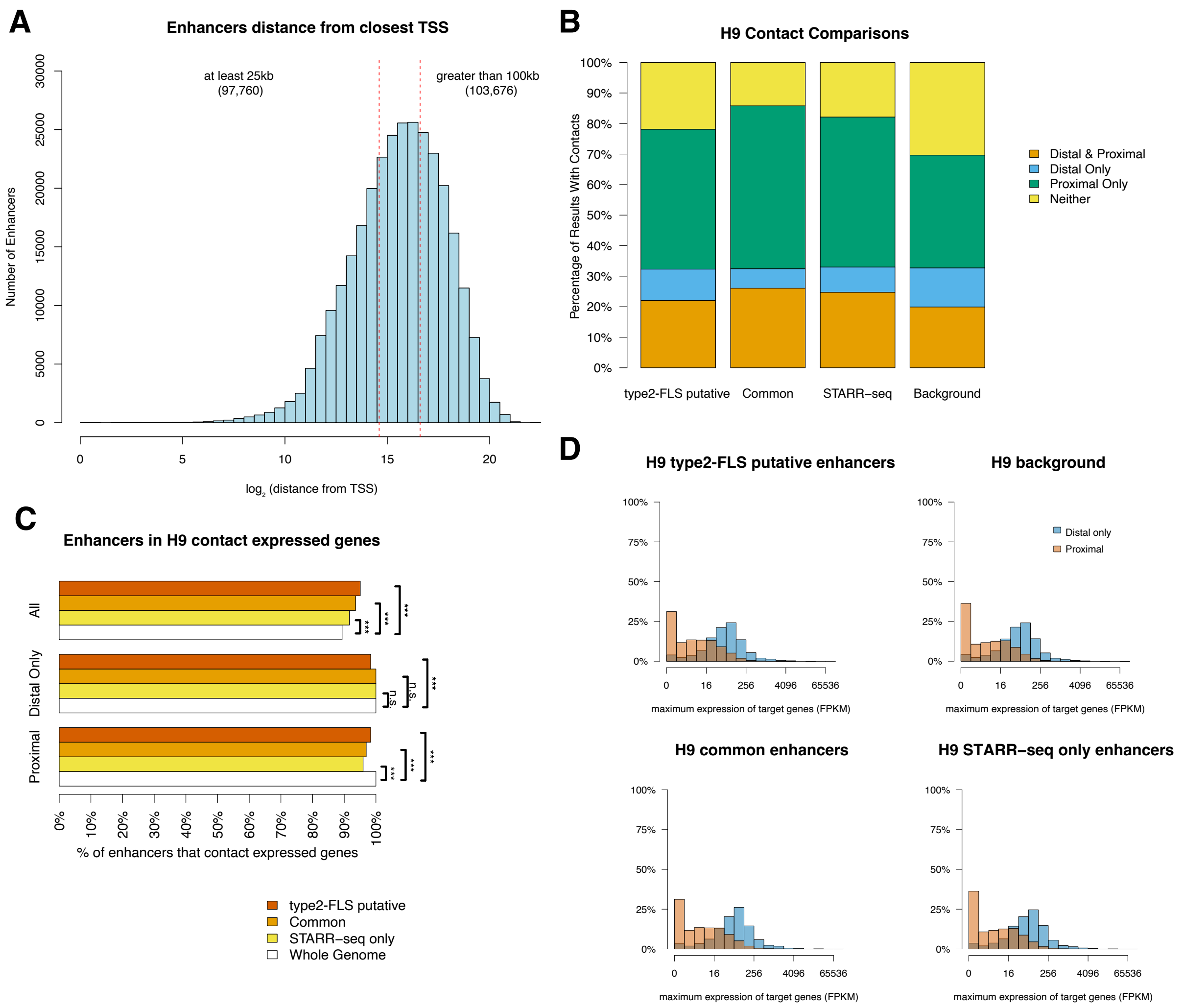


**Figure S5:** *Characterisation of proximal and distal enhancers in human ES cells. (A)* The distribution of the distance between the type2-FLS annotated enhancers that are intronic and intergenic and the nearest promoter. *(B)* We split enhancers into: *(i)* proximal if they are located within 100 Kb of a promoter, *(ii)* distal if they are further than 100 Kb from any TSS and make 3D contact with promoters (based on Hi-C data; see Materials and Methods), *(iii)* proximal only if they do not have enriched Hi-C contacts more than 100 kb away and *(iv)* neither if they are further than 100Kb from and do not make 3D contacts with any TSS. Putative, common, and STARR-seq only enhancers have enriched 3D contacts with regions containing proximal (within 100 Kb from the enhancer) or distal (further than 100 Kb from the enhancer) promoters. *(C)* The majority of the enhancers that make 3D contacts with genes contact expressed genes (Fisher's extract test; p value: n.s. ≥ 0.05, *p value < 0.05, ** < 0.01 and *** < 0.001). *(D)* Expression (FPKM) for proximal and distal only putative enhancers on a $\log_2$ scale. In the case where promoters of multiple genes were contacted, we considered the maximum expression. There is a higher expression for genes controlled by distal-only enhancers compared to proximal ones (Mann-Whitney U test of $\log_2$ of FPKM; p value $< 2.2 \times 10^{-16}$).

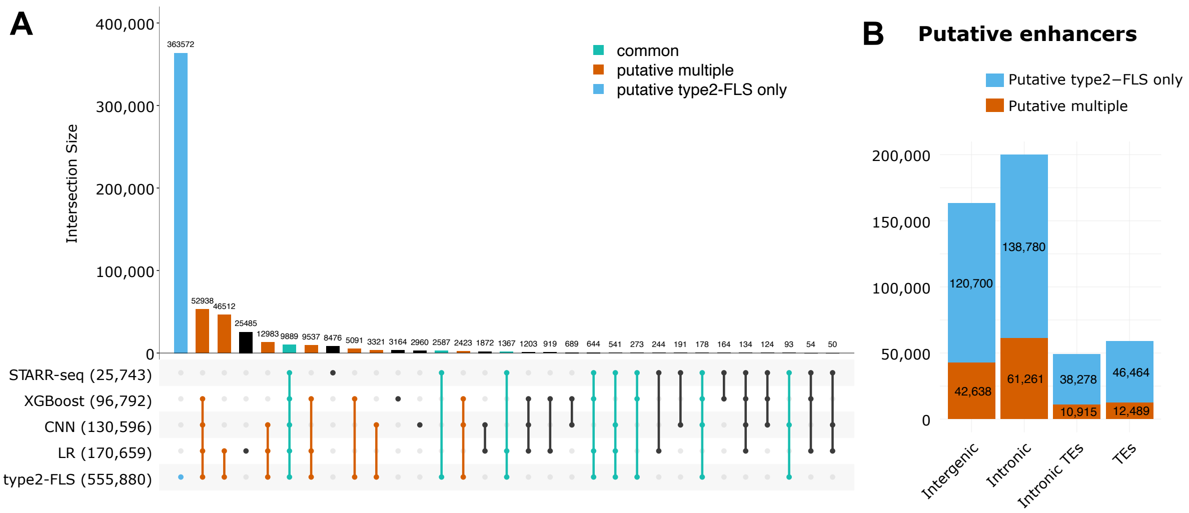


**Figure S6:** *Classification of putative enhancers*. *(A)* Upset plot with the overlap between enhancers identified by STARR-seq and the four ML/AI models (type2-FLS, CNN, LR and XGBoost) in H9 cells using 31 epigenetic features. We highlighted: *(i)* common enhancers (detected by both type2-FLS and STARR-seq), *(ii)* putative multiple (detected by type2-FLS and other ML methods but not by STARR-seq) and *(iii)* putative type2-FLS only (detected only by type2-FLS). *(B)* Number of putative multiple and putative type2-FLS only enhancers that are intergenic, intronic, intronic TEs and TEs in H9 cells.

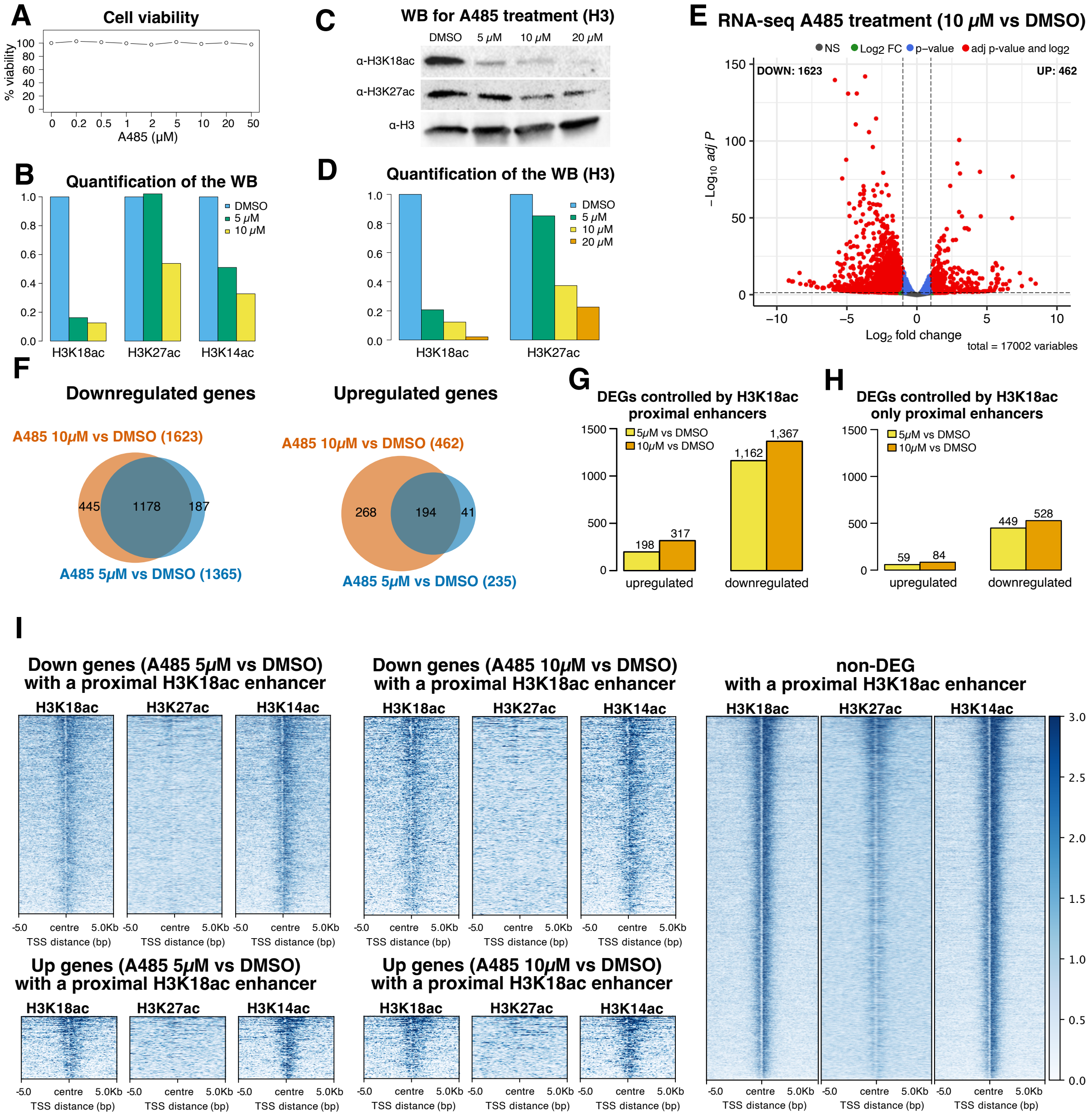


**Figure S7:** *Validation of predicted enhancers in human ES cells. (A)* Cell viability assay (Synergy HT plate reader) upon treatment with various concentrations of A485. *(B)* Quantification of Western Blot for H3K18ac, H3K27ac and H3K14ac upon A485 treatment (P300 inhibitor); see Figure 3C. We considered DMSO and 2 concentrations for A485 (5 *μM* and 10 *μM*) and used β-actin as a control. *(C)* Replication of Western Blot for H3K18ac and H3K27ac upon A485 treatment (P300 inhibitor). We considered DMSO and 3 concentrations for A485 (5 *μM,* 10 *μM* and 20 *μM*) and used H3 as a control. *(D)* Quantification of replicate Western Blots for H3K18ac and H3K27ac upon A485 treatment (P300 inhibitor); see panel *(C)*. *(E)* Volcano plot for changes in gene expression at 10 *μM* A485 compared to DMSO. *(F)* Venn diagram plotting the overlap of differentially expressed genes between A485 5 *μM* and DMSO and differentially expressed genes between A485 10 *μM* and DMSO. We considered separately (left) downregulated genes and (right) upregulated genes. *(G)* Number of differentially expressed genes that have a proximal enhancer (within 25 Kb of the TSS) that was labelled as H3K18ac positive by the type2-FLS model. We considered separately the cases of A485 5 *μM* compared to

DMSO and A485 10 *µM* compared to DMSO. *(H)* Same as *(G)* but considering only enhancers that have a medium or high level of H3K18ac and a low level of H3K27ac (as annotated by type2-FLS). *(I)* ChIP-seq signals for H3K18ac, H3K27ac and H3K14ac centred around TSS of genes: *(top left)* downregulated genes in A485 5 *µM* compared to DMSO that have an H3K18ac positive enhancer; *(top centre)* downregulated genes in A485 10 *µM* compared to DMSO that have an H3K18ac positive enhancer; *(right)* non differentially expressed genes (expressed in DMSO by at least 18 FPKM – first quartile of the expression level in DMSO for the DEGs) in both A485 5 *µM* compared to DMSO and A485 10 *µM* compared to DMSO that have an H3K18ac positive enhancer; *(bottom left)* upregulated genes in A485 5 *µM* compared to DMSO that have an H3K18ac positive enhancer; *(bottom centre)* upregulated genes in A485 10 *µM* compared to DMSO that have an H3K18ac positive enhancer.

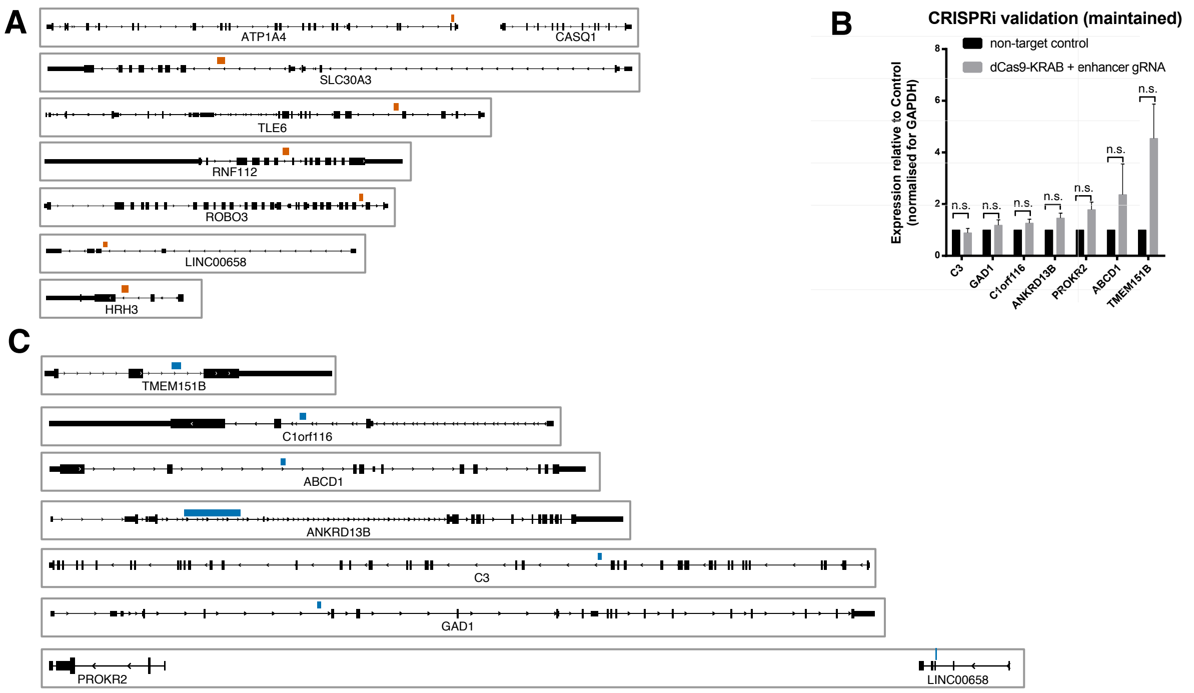


**Figure S8:** *Targeted validation of predicted enhancers in human ES cells*. *(A)* Genomic location of the enhancers targeted by CRISPRi in Figure 3E. These enhancers (red squares) are located near 8 genes, which are depicted in black. *(B)* RT-qPCR result upon CRISPRi silencing of 7 type2-FLS novel annotated enhancers. The experiment was performed in three technical replicates and three biological replicates, and a T-test was performed to evaluate if the difference is statistically significant (p value: n.s. ≥ 0.05, * < 0.05, ** < 0.01 and *** < 0.001). The enhancers plotted here are enhancers that resulted in non-statistically significant changes in the expression of the target gene. *(C)* Genomic location of the enhancers targeted by CRISPRi in in panel *(B)*. These enhancers (blue squares) are located near seven genes, which are represented in black.

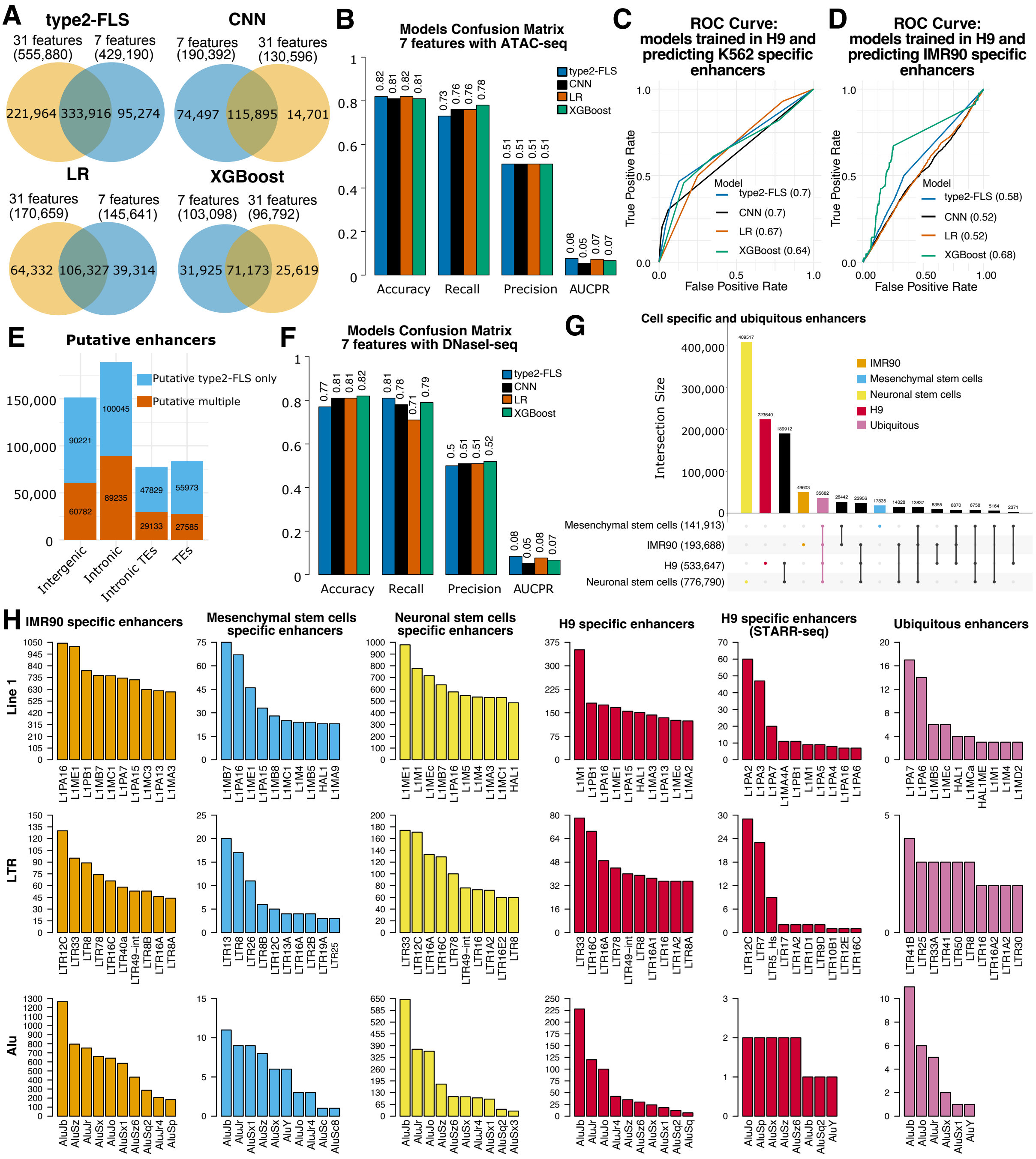


**Figure S9:** *Minimal ML/AI models and prediction of enhancers in different human cell lines. (A)* Comparison between the annotated enhancers using the 31-feature and 7-feature models in H9 cells (with ATAC-seq as a measure for DNA accessibility). We considered the case of each ML/AI method (CNN, LR, XGBoost and type2-FLS) separately. *(B)* Confusion matrix for all four ML/AI models (CNN, LR, XGBoost and type2-FLS) trained in human ES cells with 7 features, including ATAC-seq. *(C-D)* ROC curves and AUC values for four ML/AI models (CNN, LR, XGBoost and type2-FLS) trained on H9 human ES cells only on seven epigenetic features and predicting enhancers in different cell lines: *(C)* K562 and *(D)* IMR90. Training was performed on 500K bins, and the evaluation was performed on all unseen data for cell type specific enhancers. We considered true positive as the enhancers that are predicted in K562 or IMR90, and their corresponding true negative cases as active enhancers specific to the cell line where the model was trained (H9). *(E)* Number of putative multiple and putative type2-FLS only enhancers that are intergenic, intronic, intronic TEs and TEs in IMR90 cells (ATAC-seq model). *(F)* Confusion matrix for all four ML/AI models (CNN, LR, XGBoost and type2-FLS) trained in human ES cells with 7 features, including DNaseI-seq. For *(B and F)*, the values are reported on all unseen data. *(G)* Upset plot with the overlap between enhancers identified in the four cell lines (H9, Mesenchymal stem cells, Neuronal stem cells and IMR90). To annotate these enhancers, we used the 7-feature type2-FLS model using DNaseI-seq. We considered as cell-specific, enhancers in a cell line that do not overlap with enhancers in any other cell lines (using a max gap of 100 bp), while ubiquitous are the ones that are found in all four cell lines (using a max gap of 50 bp). *(H)* Number of cell-specific and ubiquitous enhancers that overlap with different subclasses of TEs. We considered separately: (top) Line 1, (middle) LTR and (bottom) Alu elements. We also considered the subset of STARR-seq enhancers in H9 cells that overlap the different TE classes.

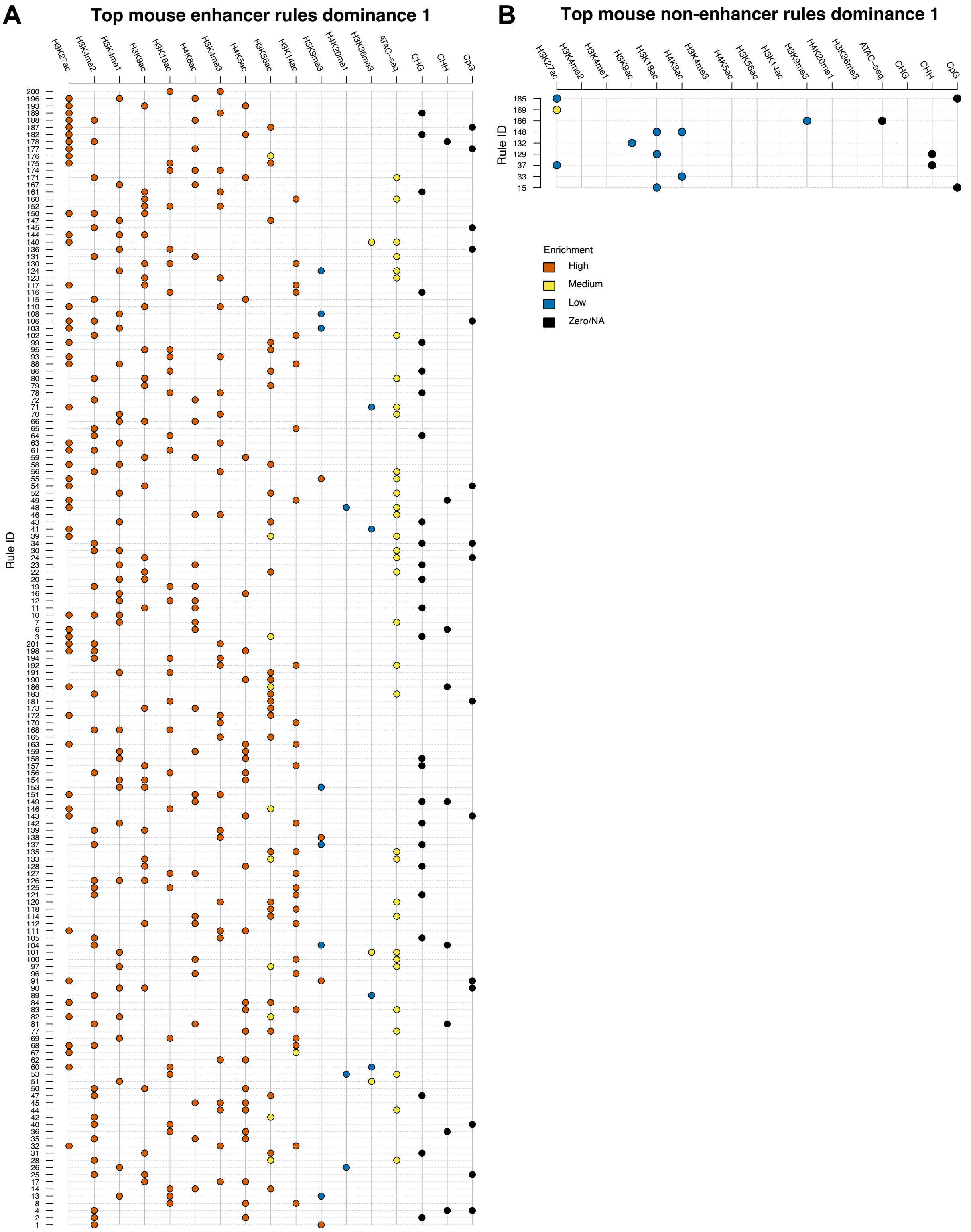


**Figure S10:** *Rules to annotate mouse enhancers.* We plot: *(A)* top mouse enhancer rules with dominance 1 and *(B)* top mouse non-enhancer rules with dominance 1. Each row represents a rule, and the colour represents the enrichment of the signal (red for high, yellow for medium, blue for low and black for zero/NA). For example, rule 200 is regions with a high level of H3K4me3 and high levels of H3K18ac are annotated as enhancers.

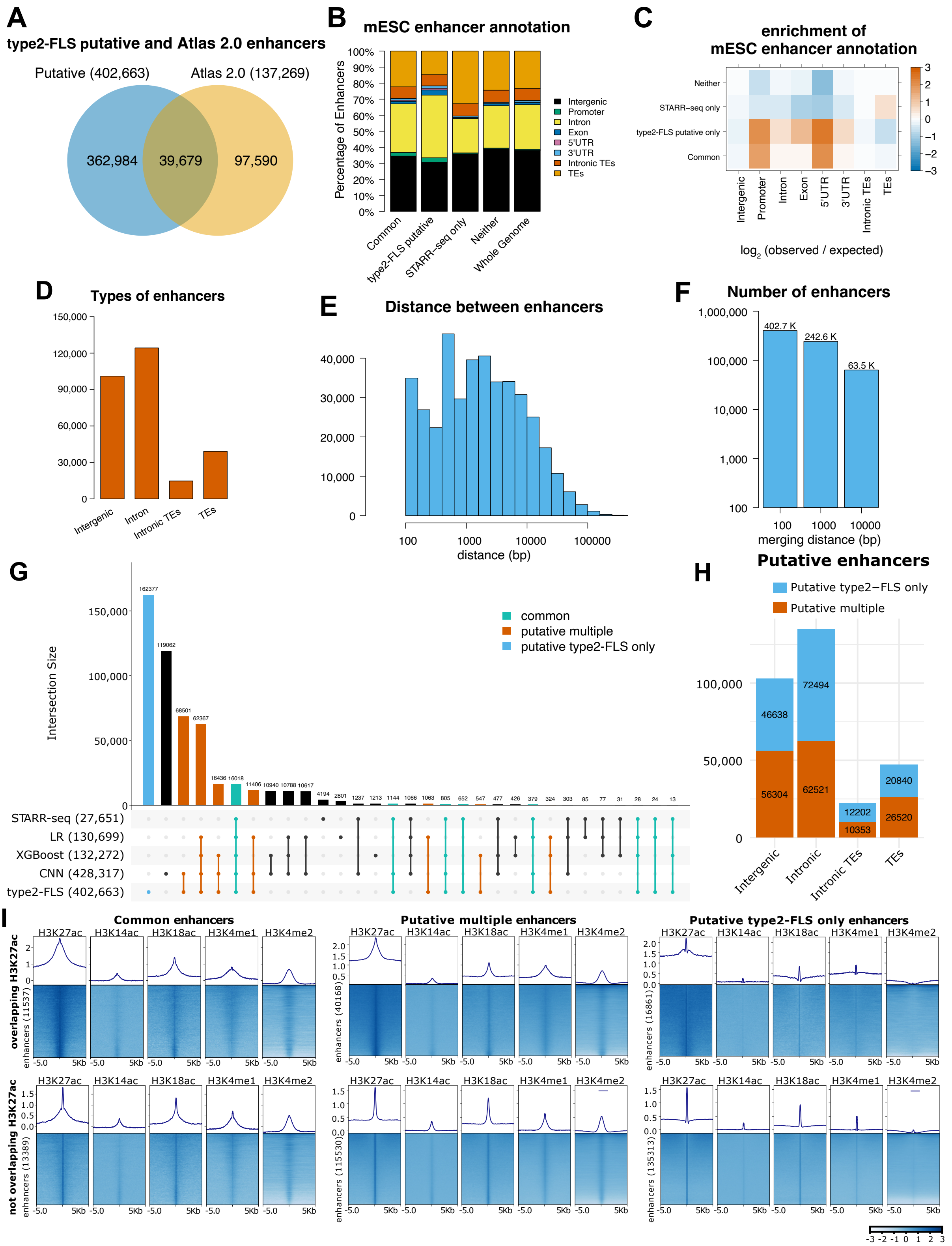


**Figure S11:** *Characterisation of mouse ES cells' putative enhancers. (A)* Overlap of putative enhancers with enhancers annotated in Enhancer Atlas 2.0. (*B)* Annotation of common enhancers (shared between type2-FLS and STARR-seq), putative enhancers (type2-FLS only), STARR-seq only enhancers and regions that are not classified as enhancers by either of the methods. We considered intergenic, promoter (up to 1 Kb upstream of TSS), intron, exon, 5'UTR, 3'UTR, intronic TEs and TEs and also plotted the whole genome distribution of the features. *(C)* $\log_2$(observed/expected) overlaps based on the whole genome distribution of the different annotations. *(D)* Barplot with the number of annotated enhancers in mESC that are located in intergenic regions, introns, intronic TEs and TEs. *(E)* Histogram of the distances to the nearest enhancer. *(F)* Number of enhancers after merging all enhancers within different distances of each other (100 bp, 1 Kb and 10 Kb). *(G)* Upset plot with the overlap between enhancers identified by STARR-seq and the four ML/AI models (type2-FLS, CNN, LR and XGBoost) in mESC using 17 epigenetic features. We highlighted: *(i)* common enhancers (detected by both type2-FLS and STARR-seq), *(ii)* putative multiple (detected by type2-FLS and other ML methods but not by STARR-seq) and *(iii)* putative type2-FLS only (detected only by type2-FLS). *(H)* Number of putative multiple and putative type2-FLS only enhancers that are intergenic, intronic, intronic TEs and TEs in H9 cells. *(I)* ChIP-seq signals for H3K18ac, H3K27ac, H3K14ac, H3K4me1 and H3K4me2 centred around mESC enhancers. We considered separately the case of*: (i)* common enhancers, *(ii)* putative multiple and *(iii)* putative type2-FLS only. We also considered separately the case of enhancers overlapping H3K27ac peaks (top panels) or not (bottom panels).

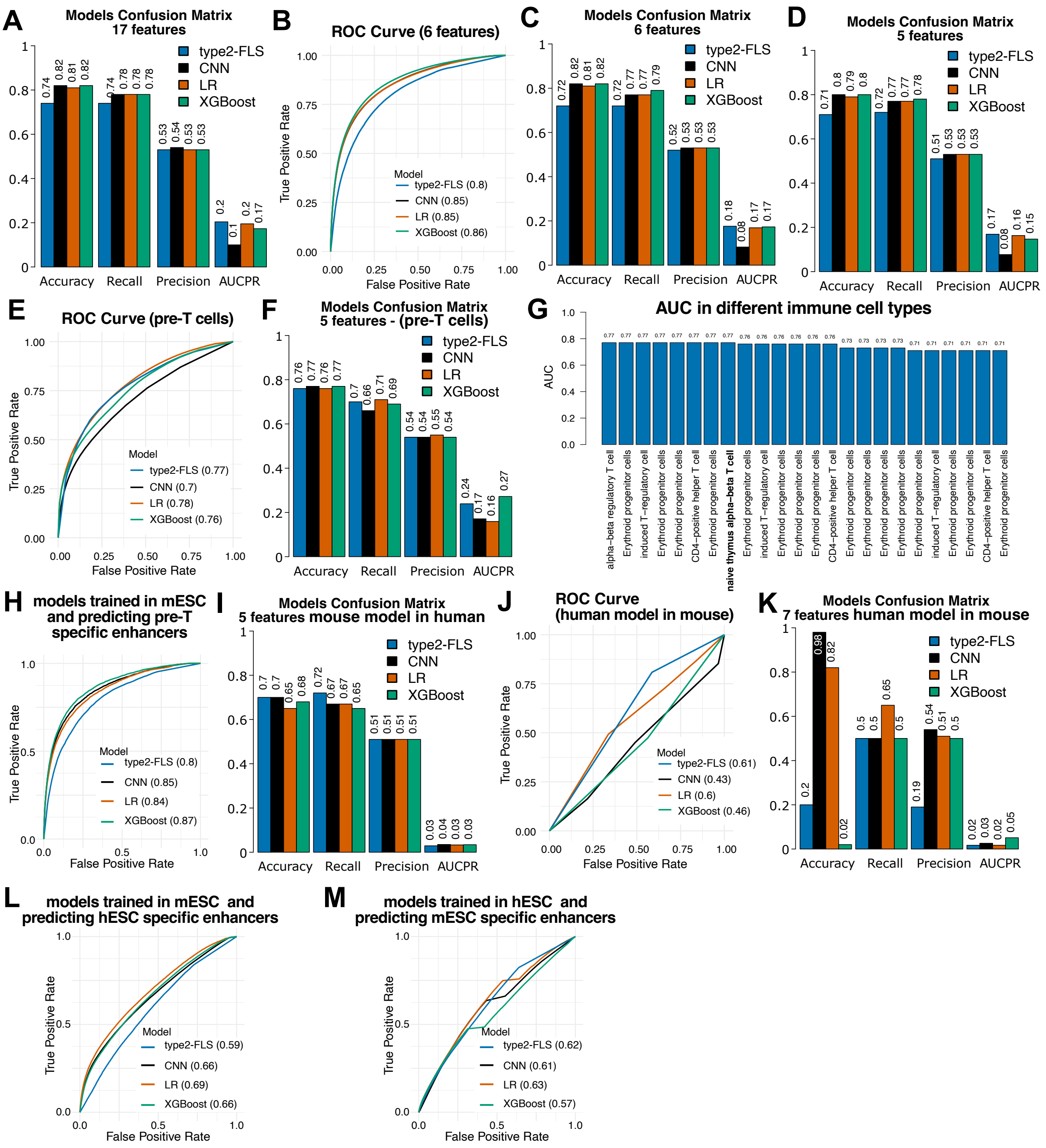


**Figure S12:** *Minimal ML/AI models and prediction of enhancers in mouse cell lines*. *(A)* Confusion matrix for the 17 features ML/AI models (type2-FLS, CNN, LR and XGBoost) trained in mESC. *(B)* ROC curve and AUC values for four ML/AI models (type2-FLS, CNN, LR and XGBoost) trained on mESC using six epigenetic features (H3K27ac, H3K4me1, H3K9me3, H3K18ac, H3K20me1 and ATAC-seq). Training was performed on 500K bins and evaluation on all unseen data. *(C)* Confusion matrix for the six features ML/AI models (type2-FLS, CNN, LR and XGBoost) trained in mESC. *(D)* Confusion matrix for the five features (H3K27ac, H3K4me1, H3K9me3, H3K18ac and ATAC-seq) ML/AI models (type2-FLS, CNN, LR and XGBoost) trained in mESC. *(E)* Evaluation of mouse minimal models in a different cell line (mouse pre-T cell line). The plot represents the ROC curve and AUC values for four ML/AI models (type2-FLS, CNN, LR and XGBoost) trained on mESC with five features and predicting enhancers in the mouse pre-T cell line on the unseen data. For enhancer annotation, we used the ENCODE annotation of enhancers in naive thymus alpha−beta T cells. *(F)* Confusion matrix for the five-feature model predicting enhancers in the mouse pre-T cell line. *(G)* AUC values for the five-feature type2-FLS model predicting enhancers in mouse pre-T cell line and using enhancer annotation from ENCODE in different mouse T cell lines. *(H)* ROC curves and AUC values for four ML/AI models (CNN, LR, XGBoost and type2-FLS) trained on mESC only on five epigenetic features and predicting enhancers in pre-T cells. Training was performed on 500K bins, and the evaluation was performed on all unseen data for cell type specific enhancers. We considered true positive as the enhancers that are predicted in pre-T cells, and their corresponding true negative cases as active enhancers specific to the cell line where the model was trained (mESC). *(I)* Confusion matrix for the five features mouse model predicting enhancers in human ES cells. *(J)* ROC curve and AUC values of the four ML/AI models (type2-FLS, CNN, LR and XGBoost) trained on human ES cells using seven epigenetic features to predict enhancers genome-wide in mESC on the unseen data. *(K)* Confusion matrix for the seven features human model predicting enhancers in mESC. All the values in this figure are reported on all unseen data. *(L-M)* ROC curves and AUC values for four ML/AI models (CNN, LR, XGBoost and type2-FLS) *(L)* trained on mESC only on five epigenetic features and predicting enhancers in hESC and *(M)* trained on hESC (H9) only on seven epigenetic features and predicting enhancers in mESC. Training was performed on 500K bins, and the evaluation was performed on all unseen data for cell type specific enhancers. We identified that 525,461 of the bins in the human genome are conserved in the mouse genome and vice versa. For *(L)*, we considered as true positives the enhancers that are only predicted in hESC by the mESC trained model (the corresponding bin in the mESC was not predicted as enhancer by the mESC trained model), and their corresponding true negative cases as mESC specific enhancers predicted by the mESC model in mESC (the corresponding bin in the hESC was not predicted as enhancer by the mESC trained model). Similarly, for *(M)*, we considered as true positives the enhancers that are only predicted in mESC by the hESC trained model (the corresponding bin in the hESC was not predicted as enhancer by the hESC trained model), and their corresponding true negative cases as hESC specific enhancers predicted by the hESC model in hESC (the corresponding bin in the mESC was not predicted as enhancer by the hESC trained model).

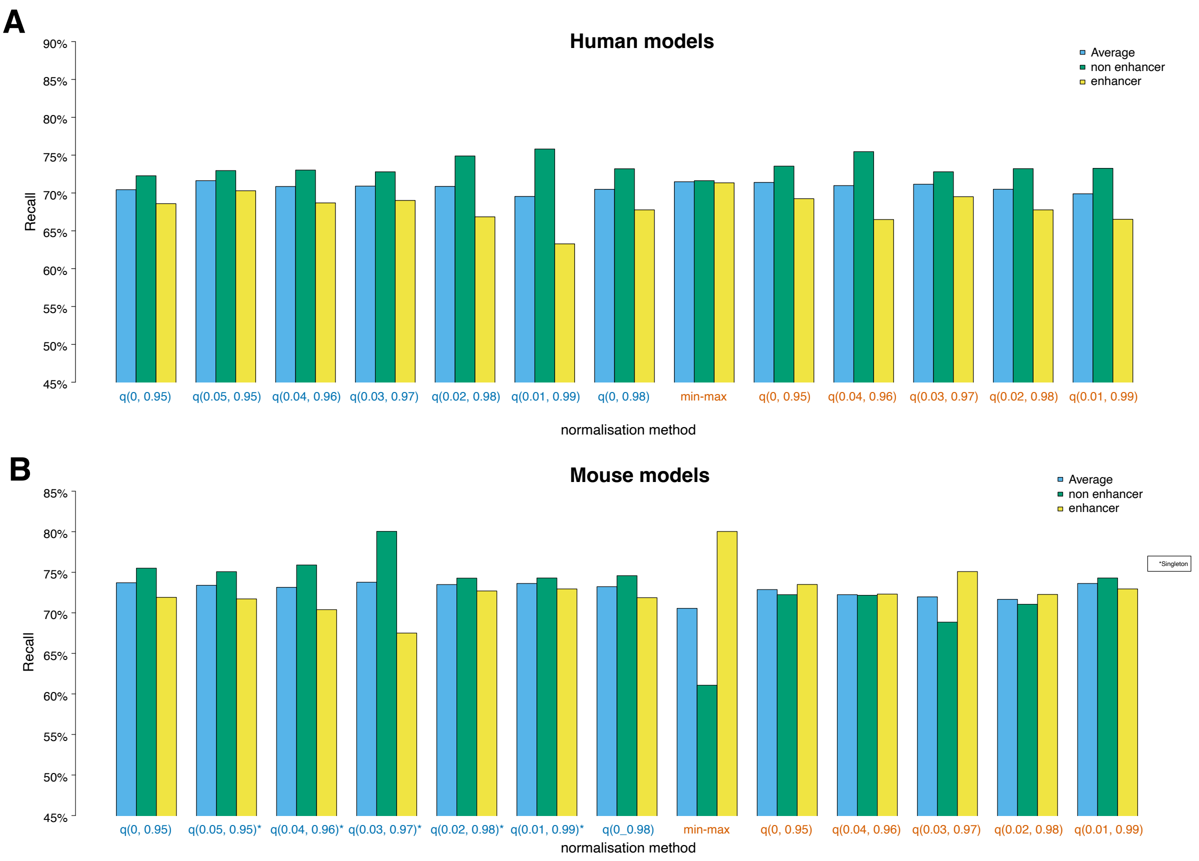


**Figure S13:** *Selection of the optimal normalisation for the data*. We trained the models on 500K bins and plotted here the recall for enhancers, non-enhancers and average recall, over the 500K holdout testing data. We optimise this for both: *(A)* human 31 features type2-FLS models and (B) mouse 17 features type2-FLS models. We considered both min-max normalisation and quantile normalisation (labelled by q with minimal value and maximum value specified for each model in the parentheses). Blue colour labels specify models where zero values were used, and red models where zero has been replaced by NA. For mouse models where we maintain the zero value, we found singletons in the data (a large number of bins with the same signal value for an epigenetic feature) identified by *.

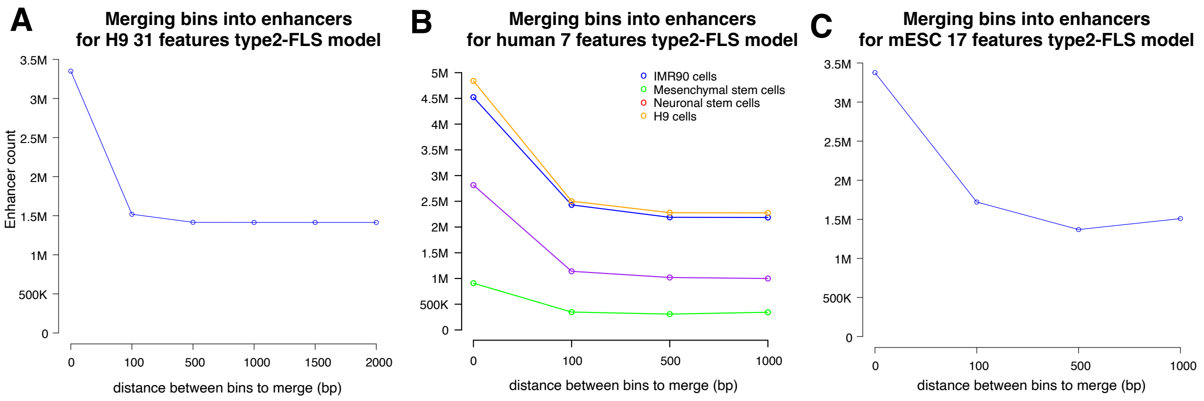


**Figure S14:** *Selection of the optimal distance to merge bins into enhancers*. We considered the case of: *(A)* 31 feature type2-FLS model in H9 cells, *(B)* 7 feature type2-FLS model in 4 human cell lines and *(C)* 17 feature type2-FLS model in mESC. We merged bins that have a probability over a specific threshold (see *Materials and Methods*) that are within a certain distance of each other (x-axis) only if the larger regions maintain the probability over the same specific threshold. Merging bins within 500 bp of each other is sufficient to minimise the number of enhancers, and using larger thresholds does not significantly reduce the number of enhancers. In panel *(C)* there is a small increase in the number of enhancers at 1 Kb, which is a small fluctuation in the total number of enhancers during the iterative merging process of the enhancers while ensuring that the entire region meets the probability criteria.
